# Revealing the Complexity of the Epicardial Secretome: Characterization and Functional Assessment of Epicardium-Derived Extracellular Vesicles and Matrix

**DOI:** 10.1101/2024.11.29.626089

**Authors:** Cláudia C. Oliveira, José Córdoba, John R. Pearson, Elizabeth Guruceaga, Ernesto Marín-Sedeño, Melissa García-Caballero, Juan Antonio Guadix, José M. Pérez-Pomares, Adrián Ruiz-Villalba

## Abstract

**Aim:** The epicardium, an epithelial layer covering the heart, plays pivotal roles in embryonic heart development and responses to cardiac damage. Epicardial secreted molecular agents are known to be involved in the regulation of these phenomena, but how this regulation occurs is poorly understood. In this study, we have investigated extracellular vesicle (EV) and extracellular matrix (ECM) components of epicardial secretions using a continuous mouse embryonic epicardial-derived cell (EPIC) line.

**Methods and results:** Epicardial-derived EVs were isolated using differential ultracentrifugation from EPIC cultured at 1% (EVs-H1%), 5% (EVs-H5%), and 21% oxygen (EVs-N). EVs protein content was determined by tandem mass tag (TMT) proteomic analysis. The results showed that epicardial-derived EVs cargo is sensitive to the oxygen level of their parenteral cells, increasing their content on glycolytic proteins. Moreover, hypoxic-derived EVs were found to both increase EPIC proliferation and affect the metabolism of Primary Human Umbilical Vein Endothelial Cells (HUVECs).

On the other hand, epicardial-derived extracellular matrix (EPIC-ECM) was characterized by subjecting decellularized EPIC to shotgun proteomics and comparing it to decellularized perinatal hearts and Matrigel®. We found that EPIC-ECM composition closely resembles that of embryonic cardiac tissue. Although the structural and basement membrane-associated proteins of EPIC-ECM were similar to those found in Matrigel®®, EPIC-ECM exhibited higher protein diversity and was a more potent inducer of HUVEC proliferation.

**Conclusion:** This work represents the first comprehensive and systematic proteomic analysis of two important components of the epicardial-derived secretome. Our experiments reveal that the epicardium responds to hypoxia by secreting EVs capable of modifying the metabolic responses of surrounding cells. Furthermore, EPIC-ECM promotes endothelial cell proliferation. These findings demonstrate the significant signaling abilities of the epicardial secretome, consistent with reports of endothelial responses following cardiac ischemic damage.

## Introduction

The epicardium is an external epithelium that covers the surface of the heart. Both cellular and molecular contributions from the epicardium are crucial for heart morphogenesis (1–4). During embryonic development, some primitive embryonic epicardial cells activate an Epithelial-to-Mesenchymal Transition (EMT), giving rise to a population of highly migratory mesenchymal cells (EPicardial-Derived Cells -EPDCs) (5–8) that can differentiate into a myriad of cardiac cell types. Epicardial epithelial cells and EPDCs secrete multiple instructive molecular signals (including retinoic acid, HGF, VEGF, BMPs, TGFβs, FGFs and IGF1 among other molecules, see (9–18). These molecules, which have mostly been regarded as paracrine signaling agents, are known to be indispensable for proper coronary vascularization (18–22) and embryonic ventricular compact myocardium growth (18, 23–26). Accordingly, loss-of-function of specific epicardial genes such as *Wt1*, *Tbx18,* or *Tcf21*, results in severe defects in both coronary and chamber myocardial wall formation, leading to midgestational death (3, 27, 28).

The role of the epicardium and EPDCs as sources of cells and paracrine signals has been extensively studied, overshadowing the critical epicardial contribution to the extracellular environment. First, the nascent ECM deposited between the myocardium and the epicardium (known as subepicardium) is a low oxygen tension milieu that modulates the EPDC differentiation, including coronary cell progenitors (27, 29–32). Furthermore, EPDCs are involved in interstitial ECM synthesis of the cardiac chamber i.e., the ECM that accumulates between cardiomyocytes, a microenvironment critical for cardiac homeostasis (2). Despite being implicated in different elements of embryonic heart development, and the onset and progression of adult heart disease, epicardial contribution to cardiac ECM remains poorly characterized.

Another neglected element of the epicardial secretome is its extracellular vesicles (EVs). EVs refers to non-replicative submicron lipid bilayer-enclosed particles that are released by virtually all cells (33). EVs are heterogeneous in size and composition, meaning that any naturally secreted particle delimited by a lipid bilayer without a functional nucleus could be considered as an EV (34, 35). Due to the difficulty in properly defining EVs diversity, an accepted nomenclature for EV relies on their size (small or medium/large EVs) (35). EVs are known to carry cellular-derived components such as nucleic acids, proteins, and lipids that are important players in cell signaling, both in health and disease (36). Interestingly, it has been shown that EV cargo composition correlates with the stimulus or biological condition triggering their formation and release (37, 38). This finding suggests the existence of intracellular selective cargo-sorting mechanisms (33, 39). Indeed, it has been reported that EVs cargo can modulate the behavior of the recipient cell. For instance, cell preconditioning to hypoxia is known to result in the release of EVs that significantly improve cardiac performance under adverse conditions (40, 41). For this reason, EVs have gained much attention in the last decade (42), having been proposed as a tool to diagnose and treat different diseases, including cardiovascular ones (43, 44).

In the present work, we have jointly characterized epicardial-secreted EVs and ECM molecules. To date, analysis of the epicardium-specific contribution to the cardiac extracellular environment has remained elusive, mostly due to technical limitations in tracing the cell-specific origin of the secretome. To overcome this limitation, we took advantage of embryonic EPIcardial-derived Cells (EPICs), a continuous EPDC line established from E11.5 embryonic murine epicardial cells that preserves various EPDC properties, including differentiation potential, active cell migration, and ECM synthesis and proteolysis (45).

Our results represent the first systematic analysis of the epicardial and EPDCs secretome, identifying potential roles for the epicardium in modulating cardiac homeostasis and responding to pathological stimuli.

## Materials and methods

### EPIC culture and extracellular vesicle isolation by ultracentrifugation

EPIC growth medium contains Dulbecco’s Modified Eagle’s Medium (DMEM, Gibco) supplemented with 10% Fetal Bovine Serum (FBS, Gibco), 100 U/mL penicillin (Sigma), 100 µg/mL streptomycin (Sigma), and 2 mM L-Glutamine (Sigma). For EVs isolation, a cell culture density of 2.7-3.8 × 10^3^ EPIC/cm^2^ was used (37 °C, 5% CO_2_). At 80% confluency, cells were cultured in EV-depleted medium (EPIC growth medium in which regular FBS is replaced by FBS ultracentrifuged at 100,000g for 5.5 h at 4°C, and filtered using a 0.2 µm-pore filter, UC-FBS). EPICs were then re-incubated at 37 °C, 5% CO_2,_ and either in 21% O_2_ atmosphere (“normoxic condition”, from here onwards EVs-N), 1% or 5% O_2_ (“hypoxic conditions”, EVs-H1% and EVs-H5%, respectively). EPIC-conditioned media was harvested every 24h for two days, and then centrifuged at 10,000 g (4 °C, 50 minutes). The supernatant was ultracentrifuged again at 100,000 g (4 °C, 70 min). The pellet was then resuspended in 0.22 µm-filtered PBS (f-PBS). EVs were washed at 100,000 g (4 °C, 70 min). EPIC-derived EVs were finally resuspended in 100 µL of f-PBS for their use in further experimentation.

In this work, we have focused on two of the most studied types of EVs: small EVs, generally considered to present a size range of 30 to 200 nm, and medium/large EVs, with a size greater than 200 nm, following the criteria of the International Society of Extracellular Vesicles (ISEV) (35).

### Transmission Electron Microscopy

Five microliters of each EV suspension were loaded onto glow-discharged Formvar/Carbon-coated 200-mesh copper grids for 15 min at room temperature (RT). Adsorbed nanoparticles were negatively stained with 1% uranyl acetate for 30 seconds. Air-dried grids were observed using a Tecnai G2 20 Twin microscope (Thermo Fisher Scientific) operating at 120 kV.

### Nanoparticle Tracking Analysis

EV particle concentration was measured with Nanosight NS300 (Malvern Instruments, UK) and Nanoparticle Tracking Analysis (NTA) software version 3.2 (Malvern Instruments). EVs were diluted at 1:1,000 and 1:2,000 in sterile water and injected into the flow cell. Samples were analyzed at identical flow rates, camera settings, and analysis detection thresholds. To confirm the cleanliness of the flow cell, measurements were also performed with sterile water. Four consecutive acquisitions (30 seconds, 25 frames/second 25°C) were recorded per sample.

### Extracellular matrix isolation

To isolate EPIC-derived extracellular matrix (EPIC-ECM), 5.3 × 10^3^ EPIC/cm^2^ were seeded in EPIC growth media that was refreshed daily until full confluency was reached. At this point, EPICs were kept in culture for three additional days to obtain an enriched ECM deposition (46, 47). NH_4_OH (20 mM) was used to induce decellularization by an osmotic shock (48). Soluble (EPIC SM) and insoluble (EPIC IM) phases were retrieved. For functional assays, urea (2M) dissolved in cold sterile PBS and filtered using 0.22-µm filter, was added to the EPIC IM fraction still attached to the plate; then the samples were scraped on ice. Plates were incubated at 4 °C for 72 hours in a rocking shaker. EPIC IM has scraped again and washed with cold sterile PBS and concentrated (250 µL) using Amicon Ultra 3 kDa cutoff centrifugal filter units (Amicon).

### Animal experimentation and embryonic heart decellularization

Animals used in this study were handled in compliance with institutional and European Union guidelines for animal care and welfare under a specific experimental procedure approved by the Ethics Committee of the University of Málaga. Mouse embryos were staged based on the presence of a vaginal plug as embryonic day 0.5 (E0.5). CD1 pregnant females were sacrificed by cervical dislocation, and embryos were isolated from the uterus and washed in PBS. Six E17.5 embryonic hearts were excised, washed with PBS, and decellularized. Briefly, the hearts were incubated at room temperature for 4 hours and 30 minutes in 20 mM NH_4_OH. The resulting embryonic cardiac mesh was then washed four times in MilliQ water and homogenized in SDS-PAGE buffer [6% (v/v) glycerol, 2% (m/v) SDS, 100 mM dithiothreitol (DTT) and 0,05 M Tris-HCl pH 6,8]. Samples were stored at −80 °C before LC-MS/MS analysis.

### Proliferation assays

To evaluate the effect of EPIC-derived EVs on either EPIC or HUVEC proliferation, 20 or 50 µg/mL of labeled EVs-N, EVs-H5%, or EVs-H1% were diluted in specific starving media added to the cells at 70% confluence and incubated for 24 h. These starving media contained 0% FBS for EPIC and 1% FBS for HUVECs (see Supplementary Material for additional information). To evaluate the effect of two different EPIC-derived ECM fractions on HUVEC proliferation, cells were seeded at 1.3-1.4 × 10^4^ cells/cm^2^ in complete medium (EGM-2) and incubated for 3 or 5 days at 37 °C and 5% CO_2_ in culture plates coated with different formulations: i) 1% gelatine (negative control), ii) 1% gelatine containing 100 µg EPIC IM, iii) 1% gelatine containing 100 µg EPIC SM, iv) 1% gelatine containing 100 µg Matrigel®(Corning), v) 1% gelatine containing 100 µg EPIC IM plus 100 µg Matrigel®, and vi) 1% gelatine containing 100 µg EPIC IM plus 100 µg EPIC SM plus 100 µg Matrigel®. Coated plates were incubated for 2 hours at 37 °C in the coating solution.

To quantify the cell number in experiments with EPIC-derived EVs or plated on EPIC IM- and EPIC SM-coated plastic dishes, cells were incubated with 5-Ethynyl-2′-deoxyuridine (EdU) (further details in the supplemental material file).

### Proteomic analysis of EPIC-derived secretome

For a quantitative analysis of the EPIC-derived EVs, Tandem Mass Tag (TMT) proteomic analysis was employed (n=3 per condition). Proteins were diluted to 1 µg/µL and extracted using Tris (2-carboxyethyl) phosphine (TCEP) and tetraethylammonium bromide (TEAB) buffers and finally digested with trypsin. Proteins were tagged using a TMT10plex Isobaric Mass Tagging Kit (Thermo Fisher Scientific) according to the manufacturer’s instructions. The tagged EVs were analyzed using Liquid Chromatography with tandem mass spectrometry (LC-MS/MS).

For a qualitative analysis of ECM composition, label-free bottom-up mass spectrometry was used. EPIC IM (n=3), EPIC SM (n=3), decellularized E17.5 hearts (n=6), and Matrigel® (n=3) were denatured and purified for LC-MS/MS analysis. Briefly, purified samples were injected into an Easy nLC 1200 UHPLC coupled to Q Exactive HF-X Hybrid Quadrupole-Orbitrap mass spectrometer (ThermoFisher Scientific). Peptides were separated at a 20 μL/min flow rate in a 50 cm analytical column (PepMap RSLC C18, 2 µm, 100 A, 75 µm x 50 cm; Thermo Fisher Scientific).

In the case of both ECM and EV peptides, elution was performed at a constant flow of 300 nL/min. The resolution of MS survey scans was set to 120,000 m/z, while for MS/MS the resolution was set to 30,000. The ion pulverization voltage was set to 2.2 kV with an m/z window from 350 to 1,500, an isolation window of 0.7 m/z, and a dynamic exclusion of 20 sec. Software versions used for data acquisition and manipulation were Tune 2.9 and Xcalibur 4.1.31.9.

### Bioinformatics analysis and statistics

For all EVs and ECM studies, a minimum of three biological replicates were assessed per condition. One-way or two-way ANOVA tests were performed using GraphPad software version 8.0.2. A p-value < 0.05 was considered statistically significant.

For TMT10plex analyzed data, *MaxQuant* (Version 1.5.4.1) software was used to identify proteins from the RAW files together with the *SwissProt* mouse protein database (release 2019_11) supplemented with common contaminants (49, 50). Proteins that were not identified in, at least, two samples of each experimental condition, would not be considered for the statistical analysis. Multivariate statistical analyses (principal component analysis – PCA - and agglomerative hierarchical cluster analysis) were performed using R/Bioconductor (51). For statistical analysis, a limma test was employed in which proteins with a p-value < 0.05 were considered differentially expressed (52). Then, EVs proteins found to be accumulated were associated with Gene Ontology (GO) terms using *clusters_to_enrichments.*R script from ExpHunterSuite (53, 54).

For label-free LC-MS/MS data, *Sequest HT* was used. Exploratory protein identification was pursued by identifying protein presence or absence in two or more biological samples with three replicates for EPIC IM, EPIC SM and Matrigel® and six replicates for E17.5 decellularized hearts. Proteins were considered part of an experimental condition if they were present in at least two replicates. The false discovery rate (FDR) for consecutive protein and peptide assignments was determined using the *Percolator* software package. GO analysis was performed at a qualitative level (presence or absence). Lists of proteins were also associated with GO terms using *clusters_to _enrichments*.R script. This script uses over-representation analysis (ORA) that selects a group of significant proteins and performs a Fisher’s exact test for each GO term. Fisher’s exact test P-value were corrected with the Benjamini-Hochberg procedure (54) the latest version of the code can be found at https://github.com/seoanezonjic/ExpHunterSuite). GO terms with adjusted P value higher than 0.05 were discarded.

Additional information on the materials and methods used in this study are available in the Supplementary Material.

## Results

### Hypoxia affects the heterogeneity of epicardial-derived extracellular vesicles

To evaluate the effect of hypoxic conditions on epicardial-derived extracellular vesicles (EPIC-EVs), we first compared the morphology of EVs isolated from EPICs cultured under different oxygen levels (Figure 1A). TEM analysis of EVs-N (Figure 1A i), EVs-H5% (Figure 1A ii) and EVs-H1% (Figure 1A iii) did not reveal significant structural differences. EVs showed a trend to equally collapse in all samples, as previously described for dehydrated EVs (55).

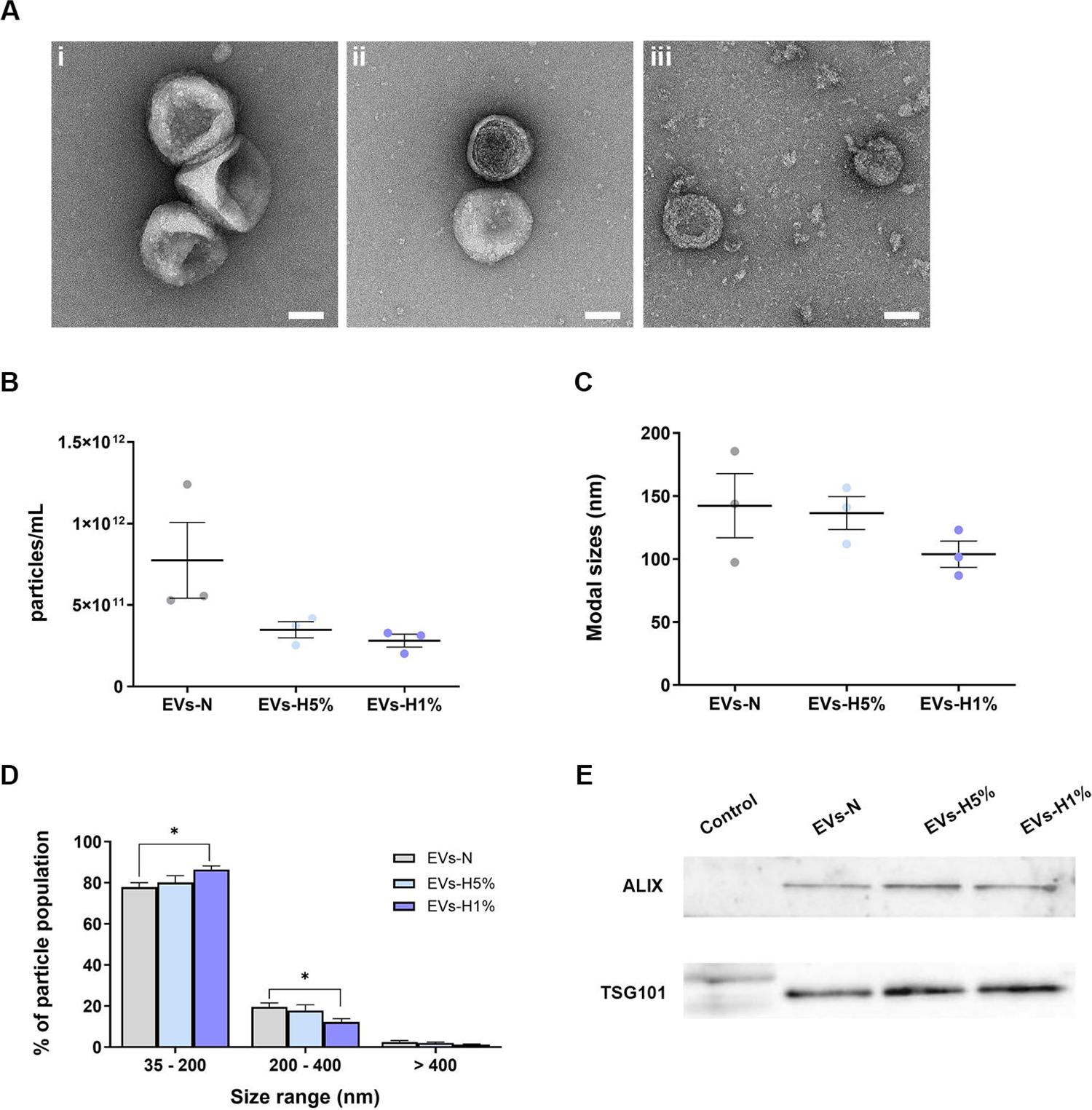

Next, we compared EVs size and concentration using Nanoparticle Tracking Analysis (NTA) (Figure 1B-D). EVs-N, EVs-H5%, and EVs-H1% showed a reduction in particle concentration with reduced oxygen concentration (Figure 1B). A similar trend was also observed in overall modal particle size (Figure 1C). We then analyzed the distribution of the particle population between small (35-200 nm), medium (200-400 nm), and large size (>400 nm) in each condition. EVs-H1% samples showed a significant increase in the proportion of small EVs and a significant decrease in medium sized EVs in comparison with EVs-N (Figure 1D). Small EV enrichment was confirmed by western blot analysis, detecting proteins typically associated with this type of EVs, such as ALIX and TSG101 (Figure 1E). Sample loading accuracy was assessed by Ponceau Red staining (Figure S1).

### Hypoxic epicardial-derived extracellular vesicles are enriched in glycolysis/glucogenesis-related proteins

To characterize and quantify potential differences between EVs isolated from EPIC cultured in hypoxic vs normoxic conditions at the proteomic level, we performed a TMT10plex study. A principal component analysis (PCA) was performed to identify similarities and differences between EVs samples (Figure 2A). We found that EVs-N and EVs-H1% proteomes were clearly separated in the PCA, whereas the EVs-H5% showed overlap with both sample groups. Next, we built a hierarchical clustered heatmap using the proteins that significantly differ between EVs-H5% or EVs-H1% and EVs-N (adjusted p-value < 0.05) (Figure 2B). This heatmap revealed 32 overrepresented proteins in EVs-H1% relative to EVs-N, 15 overrepresented proteins in both EVs-N and EV-H5% relative to EVs-H1%, and 20 proteins that decreased as the hypoxia increases.

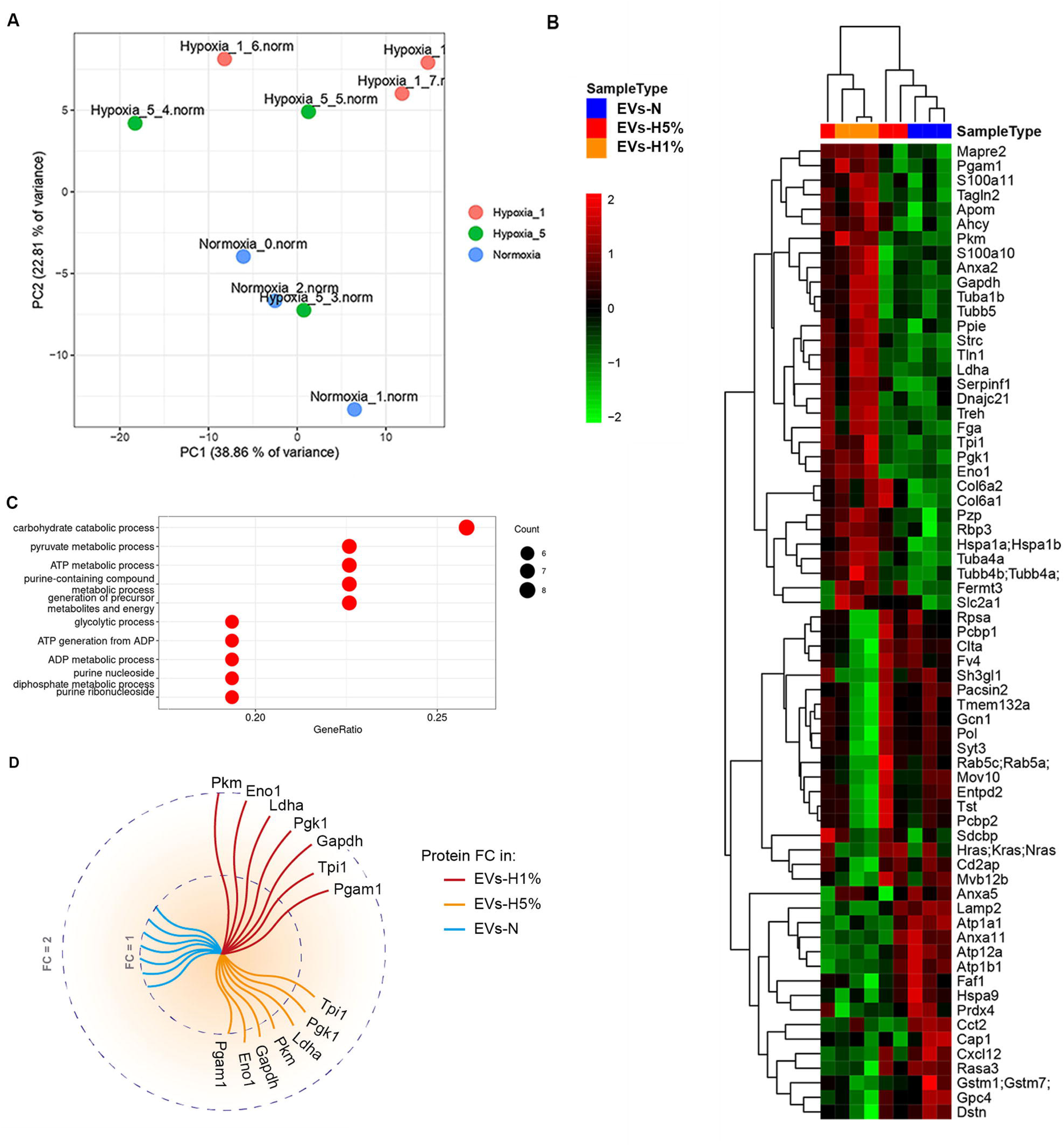

To identify biologically relevant differences related to hypoxia, we performed a gene ontology (GO) analysis of the up-regulated proteins present in EVs-H1% in comparison with EVs-N (pvalue < 0.05) (Figure 2C). Interestingly, glycolysis-related biological processes were identified among terms most significantly upregulated. Of these, GOs such as carbohydrate catabolic process (GO:0016052), pyruvate metabolic process (GO:0006090), glycolytic process (GO:0006096), ATP generation from ADP (GO:0006757), and ADP metabolic process (GO:0046031) were the most significant ones (adjusted pvalues < 1.09 × 10^-9^). Accordingly, seven glycolytic metabolism-related enzymes showed an increase in their abundancy in EVs-H1% as compared to EVs-N: phosphoglycerate mutase 1 (PGAM1; fold change (FC)= 1.47), pyruvate kinase muscle isozyme (PKM; FC = 1.95), glyceraldehyde-3-phosphate dehydrogenase (GAPDH; FC = 1.70), L-lactate dehydrogenase A chain (LDHA; 1.80), triosephosphate isomerase (TPI1; FC = 1.54), phosphoglycerate kinase 1 (PGK1; FC = 1.74) and α-enolase (ENO1; FC = 1.89). When analyzing the fold change of the same set of proteins from EVs-H5%, the FC related to EVs-N ranges from 1.04 to 1.2 (TPI1 FC = 1.20; LDHA and PGK1 FC = 1.17; PKM FC = 1.15; GAPDH FC = 1.11; ENO1 FC = 1.06; PGAM1 FC = 1.04) (Figure 2D). Altogether, these data suggest that the protein composition of EPDC-derived EVs is remarkably sensitive to the oxygen available in their milieu and showed a significant enrichment in the content in glycolytic enzymes related to the increase in the hypoxia grade to which the parenteral cells were subjected.

### Extracellular vesicles derived from hypoxic epicardial-derived cells exert both autocrine and paracrine effects in surrounding cells

Based on the results of our TMT10plex analysis, we hypothesized that epicardial-derived cells could affect the glycolytic function of surrounding cells via EVs. To assess this, we studied the potential autocrine effect of EPIC-EV on the EPIC line itself. In parallel, we tested the paracrine effect of these EVs on human umbilical cord-derived endothelial cells (HUVECs).

To evaluate the internalization of EPIC-EVs, samples were stained with lipophilic dyes just before incubating with different cell types. On one hand, EVs-N were stained with DiI (EVs-DiI; Figure S2) and added to cultured EPICs. The internalization of EPIC EVs was confirmed by the counterstaining of acidic organelles of EPIC with LysoTracker^TM^ (Figure 3A). On the other hand, EPIC-EV internalization by HUVECs was also confirmed after staining EVs with PKH26 and using LysoTracker^TM^ to counterstain HUVEC lysosomes (Figure 3B). Then, to evaluate potential changes in the biology of target cells, cell proliferation was analyzed in both EPICs and HUVECs after incubation with different concentrations (20 or 50 ug/mL) and types of EVs (EVs-N, EVs-H5% or EVs-H1%) (Figure 3C-D). No significant differences in cell proliferation were found between the basal condition and EPIC incubated with EVs-N or 20 ug/mL of EVs-H5%. However, the number of EdU^+^ EPIC was significantly increased after incubation with 50 ug/mL EVs-H5%, and both concentrations of EVs-H1% (Figure 3C). On the other hand, HUVEC proliferation did not significantly change after incubation with each EVs type as compared to its basal condition (Figure 3D).

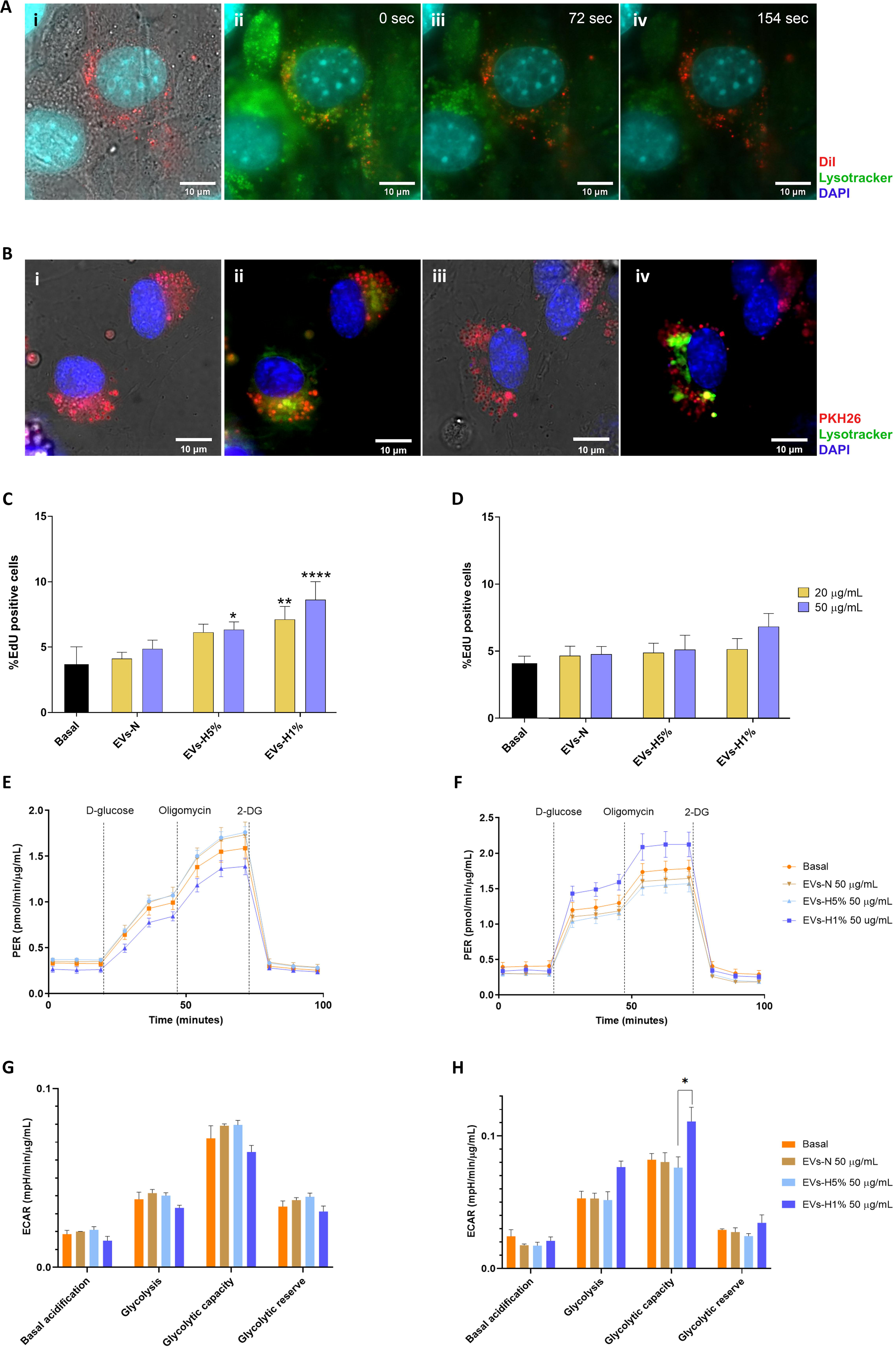

Next, we evaluated the capacity of EPIC-derived EVs as potential carriers of metabolic modulators to induce a glycolytic response in the receiving cells using a Seahorse Glycolysis Stress Test. For doing that, EPICs or HUVECs were incubated with 50 μg/mL of EVs-N, EVs-H5%, or EVs-H1%. First, none of the different populations of EVs induced significant changes to the glycolytic behavior of EPIC, but EVs-N andEVs-H5% tended to increase both behaviors. However, EPIC incubated with EVs-H1% tended to drastically reduce their glycolysis and glycolytic capacity (Figure 3E and G). Interestingly, HUVECs incubated with EVs-N and EVs-H5% tended to reduce their glycolysis and glycolytic capacity. But in the presence of EVs-H1%, HUVECS drastically increased their glycolytic capacity in comparison with HUVECs incubated with EVs-H5% (Figure 3F and H). This data suggests that EPIC-EVs can modify the metabolism of the receiving cells.

### Soluble and insoluble protein fractions can be isolated from epicardial-secreted extracellular matrix

To analyze another potentially important element of the epicardial secretome, we explored the composition of the epicardial-derived extracellular matrix (EPIC ECM). This procedure resulted in the differential isolation of two ECM fractions, one soluble (EPIC SM) and the other insoluble (EPIC IM). When analyzed by scanning electron microscopy, the EPIC IM fraction displayed a rough surface and a clear irregular organization in comparison with EPICs, that showed fibers assembled to form a mesh-like organization (Figure 4A). Routine immunohistochemical analysis of both EPIC ECM fractions revealed they are enriched in basement membrane proteins and contained low (EPIC SM) or negligible (EPIC IM) amounts of intracellular proteins (Figure S3).

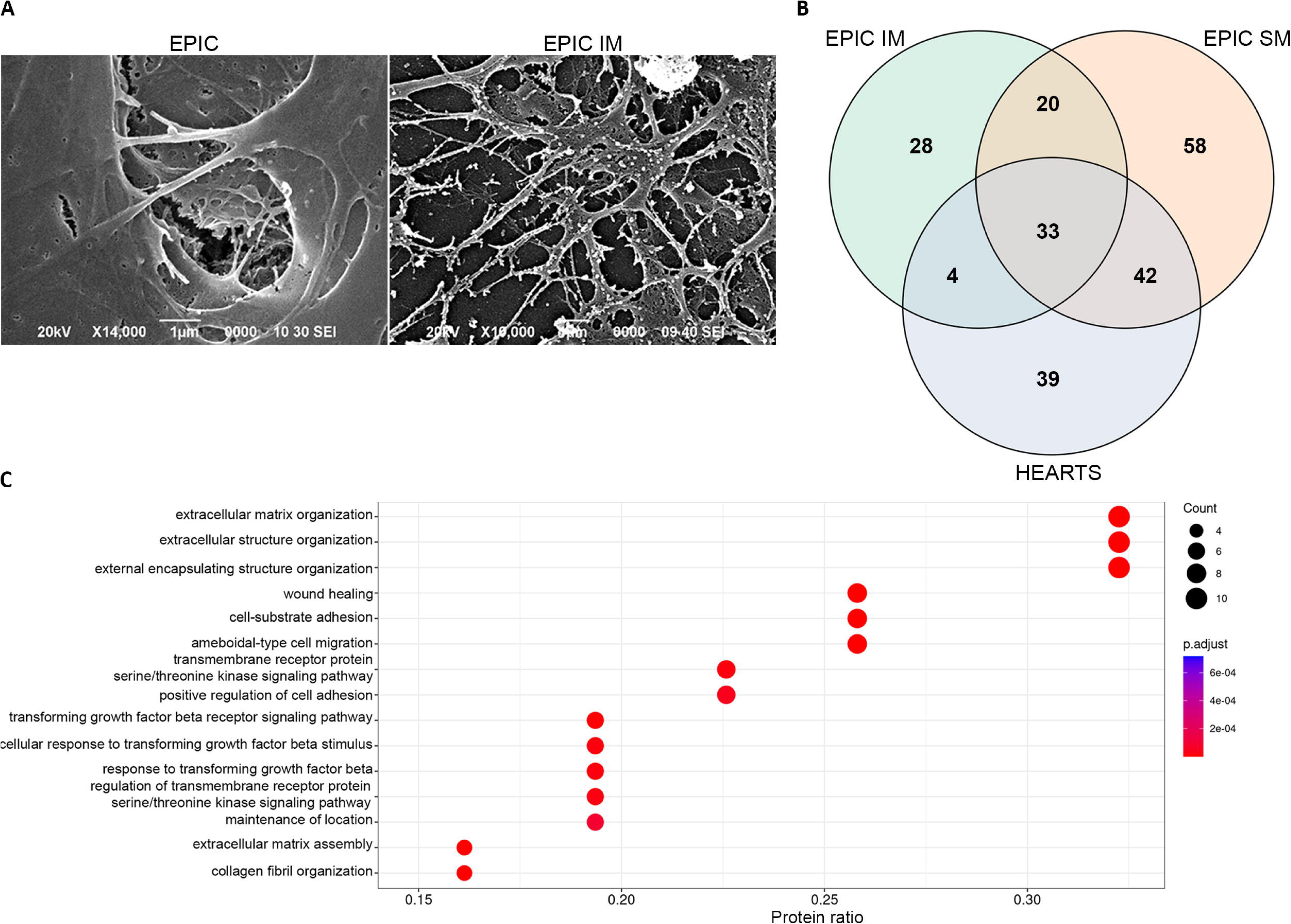

To determine their extracellular protein composition, both ECM fractions were submitted to shotgun proteomics. Based on GO annotations, 24% of EPIC IM and 9% of EPIC SM identified proteins were identified as “extracellular”. A total of 53 extracellular proteins were found to be common between the two fractions. This includes several cardiac-enriched ECM proteins such as hyaluronan, proteoglycans, elastin, fibrillin, and tenascin (Table S1). As expected, all these proteins are associated with biological processes related to the ECM such as extracellular matrix organization (GO:0030198), extracellular structure organization (GO:0043062), wound healing (GO:0042060), collagen fibril organization (GO:0030199), cell-substrate adhesion (GO:0031589), or extracellular matrix assembly (GO:0085029) (adjust p-value < 1 × 10^-7^) (Figure S4).

To define their unique protein signature, we identified the subset of proteins present only in each EPIC-ECM fraction. In EPIC IM, 32 unique proteins were identified, including more than ten types of collagen and other proteins related to cardiac ECM such as AGRN, FGFR2, or SPARC. Other EPIC IM-specific proteins were PDGF, SERPINF1, CTGF, ADAMTS1, SEMA3E, SEMA3B, and NPNT (Table S2). For EPIC SM, 100 unique proteins were found, such as CXCL12, PDGFB, TGFB2, or WNT7A (Table S2), all directly related to signaling pathways associated with cardiac ECM development and repair. The protein signature of the two fractions can be associated with similar GO terms related to angiogenesis, migration, proliferation, or cell adhesion (Figure S5). However, proteins exclusive from the EPIC IM fraction showed a better association with these GO terms (0.2 protein ratio) than the ones from the EPIC SM fraction (0.11-0.14 protein ratio).

### Comparative analysis of the epicardial-derived extracellular matrix composition

To evaluate the potential significance of the epicardial contribution to cardiac ECM, we compared both EPIC-derived ECM fractions with the ECM isolated from a pool of decellularized murine embryonic hearts (DMEH). Shotgun proteomics identified 33 extracellular proteins in common between the three samples (28%), such as FN1, COL1A1, or COL1A2 (Figure 4B, table S3). To better characterize the similarities between EPIC ECM and DMEH proteomes, a GO term analysis was performed (Figure 4C). As expected, these 33 proteins were associated with biological processes related to ECM organization, wound healing, cellular adhesion and migration, and molecular signaling. In addition, 40% of the detected extracellular proteins are common between DMEH and each one of the fractions. Only 34% were unique to the DMEH fraction, showing a high degree of similarity between EPIC ECM and DMEH.

To understand the specificity of EPIC-derived ECM protein composition, we compared it with Matrigel®, a widely used commercial ECM for *in vitro* cell growth studies. Using label-free LC-MS/MS, 314 proteins were identified in Matrigel® samples, but only 89 of them were annotated as “extracellular”. Out of these 89 extracellular proteins, 46% were found to be exclusive for Matrigel® and associated with GO terms related to the negative regulation of different enzymatic activities, such as hydrolase (adjust p-value = 5.74 × 10^-12^), peptidase (adjust p-value = 9.95 × 10^-13^) or proteolysis (GO:0045861; adjust p-value = 2.25 × 10^-11^) (Figure S6A, B). In contrast, 23 extracellular proteins were identified in Matrigel®, EPIC IM and EPIC SM samples (Figure S6A). Most of these common proteins belong to laminin (LAM) and heat shot protein (HSP) families and are associated with GO terms related to extracellular functions, such as extracellular matrix organization (FDR of 1.55 × 10^-5^), substrate adhesion-dependent cell spreading (FDR of 1.56 × 10^-5^), and regulation of localization (FDR of 1.56 × 10^-5^) (Figure S6C). Taken together, our results indicate that the proteins found in the EPIC-ECM fractions are substantially different from those identified in Matrigel®.

To further evaluate the specificity of EPIC IM and SM fractions, all our generated datasets (EPIC IM, EPIC SM, DMEH, Matrigel®) were compared (Table 1). The first analysis we carried out revealed that only 17 extracellular proteins, associated with LAM and HSP families, were shared between the four datasets (Figure S7A and Table S4). A GO term analysis from these 17 proteins identified a set of biological processes related to cell attachment, flattening, and differentiation, such as substrate adhesion-dependent cell spreading (FDR of 3.18 × 10^-5^), cell morphogenesis involved in differentiation (FDR of 3.18 × 10^-5^) and biological adhesion (FDR of 8.20 ×10^-5^) (Figure S7B). To unravel the specificity of the composition of each sample, the relative abundances of extracellular proteins in the total protein pool per condition were calculated (Table S5). In EPIC IM, serine protease HTRA1 represented 15.21% of total protein, followed by CXCL12 (5.26%), FN1 (3.36%), EMILIN-1 (3.01%) and LAMA1 (2.76%). On the other hand, EPIC SM was highly enriched in FN1, showing an abundance reaching 40.43% of the total proteins identified, followed by MYH9 (19.76%), endoplasmin (HSP90B1) (2.92%), ANXA2 (2.55), and HSPA5 (1.94%). For Matrigel®, three laminins were the most abundant proteins: LAMA1 (33.09%), LAMB1 (22.30%), and LAMC1 (19.43%) (Table 1). These results were validated by western blot for both FN-1 and LAMA1 (Figure S8). On one hand, FN-1 detection was stronger in the EPIC SM than in the EPIC IM, Matrigel® or DMEH (Figure S8A). On the other hand, Matrigel® presented the strongest signal for LAMA1, as expected. EPIC SM also showed signal at the top of the membrane, whereas no bands were detected in EPIC IM, EPIC lysate and DMEH (Figure S8B).

Taken together, our analysis shows that EPIC secretes ECM proteins phenotypically related to an embryonic murine heart, whereas it distances from non-cardiac and non-embryonic ECM-derived tissues, such as Matrigel®. EPIC-ECM closest resemblance to Matrigel® relates to main ECM roles such as structural and organizational functions.

### Epicardial-derived extracellular matrix induces the proliferation of endothelial cells

To validate whether the differences in protein composition extends to a functional specificity, we performed a functional study to evaluate the potential effect of EPIC-ECM on HUVEC growth. HUVECs were cultured on wells coated with different substrates, combining gelatine with EPIC IM, EPIC SM, and/or Matrigel® (Figure 5). The HUVEC proliferative rate, measured as a percentage of EdU-positive cells, was evaluated after three (Figure 5A) and five days (Figure 5B) of culture.

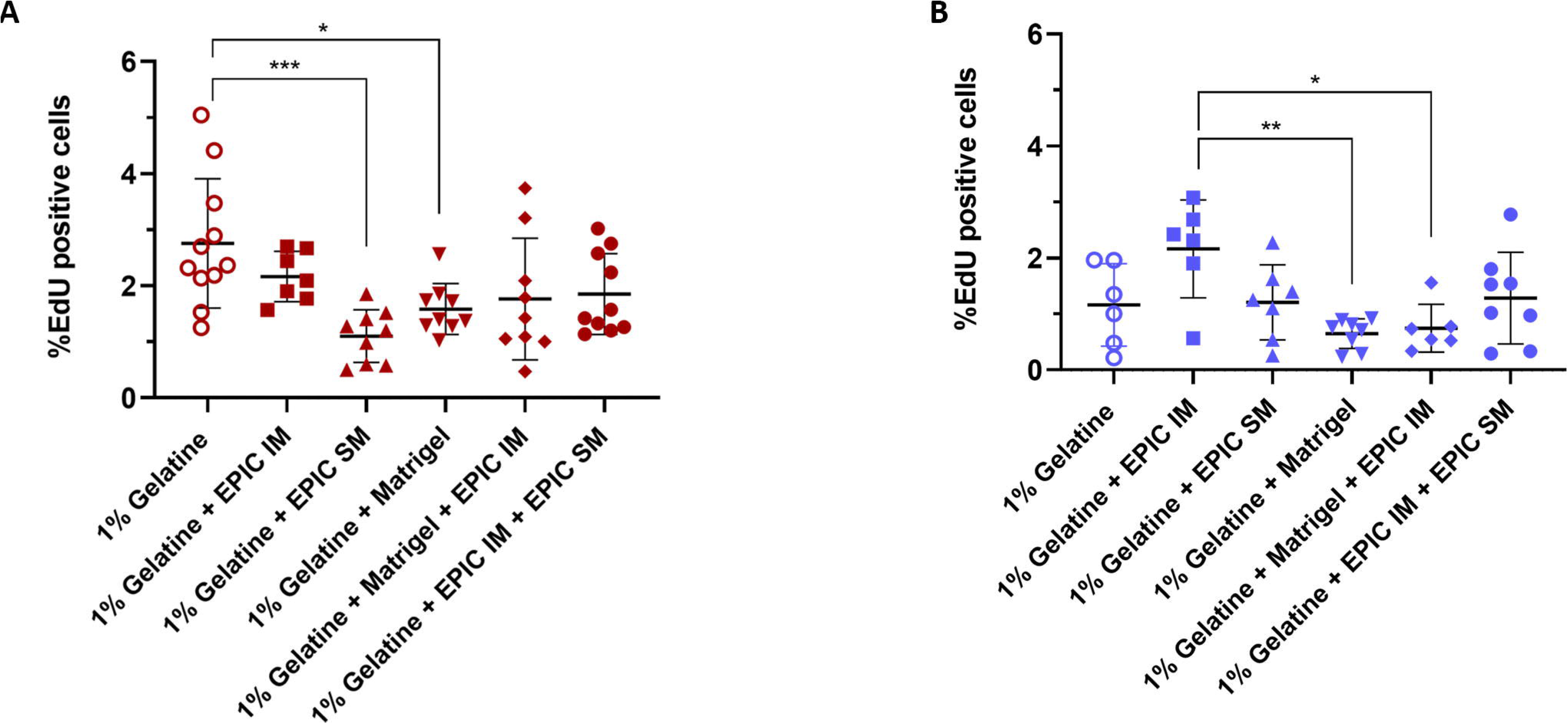

After three days, the proliferation rate of HUVECs cultured on gelatine plus EPIC SM or Matrigel® was significantly lower than in wells coated only with gelatine (Figure 5A). However, after five days, HUVECs plated onto both types of substrates containing Matrigel® showed a lower percentage of EdU-positive cells versus cells cultured only on the EPIC IM fraction (Figure 5B). These results indicate that EPIC IM and Matrigel® have functionally different effects on HUVEC growth during the 5-day assay.

## Discussion

Several reports have demonstrated the importance of the epicardium and EPDCs during the formation, growth, and maturation of the heart. The function of this cell lineage has been attributed to its role as the source of coronary vascular and perivascular cells, and cardiac fibroblasts (45, 56–60). As pioneers in the colonization of the nascent cardiac interstitium, a complex space that is crucial to cardiac homeostasis (2), EPDCs actively participate in the conditioning of the extracellular cardiac microenvironment during both development and disease through the secretion of multiple molecules into the extracellular milieu (61, 62). However, despite its enormous relevance to the pathophysiology of many cardiac diseases, there have been few studies into the EPDC-derived secretome. This is due to technical challenges related to the low number of native epicardial and EPDCs present in the embryo under normal conditions. To circumvent these limitations, we took advantage of EPIC, an immortal cell line derived from mouse embryonic epicardium that has been shown to recapitulate an important part of embryonic epicardial biology (45). In this report, two fractions of the epicardial secretome were studied: EVs and the ECM.

EVs are relevant to cell-to-cell communication (63–65) and can reflect changes in the physiological status of the parental cells, such as changes in oxygen availability (66–69). Since the embryonic heart develops under low oxygen tension conditions (31, 70), and hypoxia modulates the differentiation of EPDCs (27, 29, 30, 32), the effect of hypoxia over epicardial-derived EVs was investigated. First, we found EPIC-derived EVs to be morphologically similar to those released by other cell types (71, 72). However, hypoxic EPICs secreted smaller-sized EVs than normoxic ones, as reported for other cell types in different biological contexts such as cancer (73–75) or liver fibrosis (76). Moreover, the protein signature of EPIC-derived EVs gradually shifts towards a composition enriched in glycolysis/gluconeogenesis enzymes as oxygen levels in EPIC cultures decrease. Interestingly, this enrichment was accentuated in EVs-H1% versus EVs-H5%, indicating that the cargo of EPDC-derived EVs is remarkably sensitive to the oxygen available in their milieu.

Our study also shows that the highest concentration of EVs-H5% and both of EVs-H1% enhanced EPIC proliferation, pointing to a likely autocrine role for epicardial EVs. Unexpectedly, this inducement is not directly related to a glycolytic response, as shown by our Sea-Horse data. Our results contrast with several works where enzymes as PKM2 or PGAM1, significantly enriched in EVs isolated from hypoxic EPIC, promote proliferation in other cellular contexts, such as cancer (77, 78). However, it has been described that the cardiac overexpression of PGAM2 affects the cell metabolism at different level, including the mitochondrial respiration rate and the generation of reactive oxygen species (79). Based on that, epicardial cells may have the capacity to respond to high energy demands without drastically changing their metabolism, as a pre-adaptation to their cardiac hypoxic niche during development (31). Moreover, and considering the immortalization process suffered during their generation (45), it is possible that specifically EPIC metabolism could be more dependent on amino acids such as L-Glutamine, an essential component of their culture medium, than on glucose, as described in other immortalized cell lines (80).

On the other hand, EVs-H1% induced a drastic but not significant change in the glycolysis response and glycolytic capacity of the HUVECs, although did not enhance cell proliferation in this cell type. Considering that glycolysis is the primary energy-producing mechanism in proliferative endothelial cells (81), these results suggest that epicardial-secreted EVs may induce a glycolytic preconditioning in the targeted endothelial cells, perhaps through the incorporation of glycolytic enzymes enriched in EVs-H1% such as PKM, ENO1 or PGAM1 (82–85). However, other metabolic pathways that also affect the proliferation of endothelial cells, such as amino acid metabolism, fatty acid β-oxidation and oxidative phosphorylation, cannot be discarded (86).

Taken together, these results show that the internalization of EVs cargo triggers an altered metabolism of the receiving cells, indicating the capacity of epicardial-derived cells to affect the cellular environment in both an autocrine and a paracrine manner. In this scenario, it is important to highlight that embryonic cardiac interstitial EPDCs persist until adulthood and have been reported to play an important role in cardiac responses to pathological conditions (87). This is especially relevant in the case of ischemic events leading to a marked decrease in oxygen levels in the ventricular walls, considering the hypoxic EV-mediated response as a known mechanism that modulate cellular behavior under adverse conditions (88). In this regard, it has been recently reported that the hypoxic conditions that often follow tissue damage can induce the metabolic reprogramming of quiescent cardiac fibroblasts into a differentiated/activated phenotype. This phenotype is characterized by using anaerobic glycolysis as their main energetic source (89). If we consider that the adult cardiac fibroblasts activated after myocardial infarction are derived from the embryonic epicardium (59), it is reasonable to hypothesize that EPDC-derived EVs could act as an autocrine, endogenous mechanism of cell-to-cell communication during pathologic ventricular remodeling. In addition, the metabolic reprogramming of these anaerobic, activated cardiac fibroblasts after injury has been proposed as a potential candidate for novel therapeutic strategies to treat myocardial infarction (90). Relevant to this discussion, Villa Del Campo and colleagues reported on the miRNA contents of epicardial-derived EVs, and associated cardiomyocyte proliferation and improved cardiac contractility to this specific EV cargo (91). Although our work provides novel and relevant information on the EVs secreted by epicardial cells, the role of EPDC-derived EVs in pathological contexts needs to be further explored, including a more detailed molecular characterization of EPDC-derived EVs.

The second part of our work focuses on the extracellular matrix (ECM) derived from the epicardial cells. As previously discussed, EPDCs originally contribute to the formation of the subepicardial space, an ECM-rich environment that separates the epicardium from the myocardium, supporting early coronary vascularization (27, 61), and the formation of the ventricular interstitial space (2). Therefore, it is obvious that the molecular composition of the epicardial-derived ECM is crucial to understanding how cells respond to changes in their microenvironment (61). So far, most studies into epicardial-secreted ECM proteins are limited to a general expression mapping of a few molecules in the vicinity of the epicardium. The difficulty of tracing the specific cellular origin of secreted proteins in the embryo, and the high number of cells required for studying epicardial-derived components, have so far prevented a larger-scale analysis of the epicardial ECM secretome. To circumvent these constraints, we took advantage of the ECM synthesized by the EPIC line and completed the most systematic characterization of epicardial-derived ECM proteins to date.

Using shotgun proteomics, we found representative cardiac ECM proteins, such as hyaluronan, proteoglycans, elastin, fibrillin, and tenascin, in both the soluble and insoluble fractions of our experimental samples (EPIC SM and IM, respectively). Our results also indicate that EPIC-derived ECM contains proteins associated with both the basement membrane of the epicardium, such as laminins, FN-1 and COL4A1 and COL4A2, and the interstitial subepicardial matrix, such as COL1A1, COL1A2 and COL3A1 (61, 92, 93). Most of them have already been reported as proteins synthesized by the epicardium during development (94). Remarkably, both EPIC ECM fractions contain proteins associated with vascular growth and wound healing phenomena, in accordance with our understanding of the epicardial contribution to coronary vascular development in the developing heart (e.g. FN-1, LOX, ANXA2, FBLN1, MGP, CTHRC1) (95–101). Moreover, we demonstrate that the EPIC-secreted ECM protein profile contains similar proteins than the ECM of mouse embryonic hearts, such as LAMB1 and 2, FN-1, COL1A1 and 2, FIBL-2, HSPG and EMILIN-1 (61, 102–105). Notably, the importance of these components in cardiac morphogenesis has been demonstrated by genetic linkage to congenital heart malformations (106–108), suggesting that EPIC ECM composition is similar to the naïve epicardial-derived ECM.

Finally, we evaluated the specificity of the EPIC ECM in comparison with Matrigel®, the gold standard basement membrane-derived ECM gel. As expected, Matrigel® was enriched in proteins typically found in many basement membranes, such as laminins, NID-1, or HSPG (109). Accordingly, lower abundances of these molecules were identified in both EPIC ECM fractions and decellularized mouse embryonic hearts. However, Matrigel®’s similarity to EPIC-ECM is limited to a few protein elements, whereas a strong compositional resemblance was found between EPIC-ECM and embryonic heart ECM, both enriched in proteins associated with vasculogenesis and wound healing mechanisms. Accordingly, we confirmed differences in the proliferation of endothelial cells depending on the substrate used to grow them. When seeded on EPIC IM and EPIC SM, endothelial cells maintained their cell proliferative capacity over five days, whereas endothelial cells seeded on Matrigel®, their proliferative capacity was reduced drastically after five days of incubation. Based on our data, we hypothesized that growth effects could result from the effect of ECM-bound growth factors and chemokines released in response to the growing cells (e.g. by the partial degradation of the substrate). This interpretation is compatible with the presence of chemokines, such as CXCL12 (a.k.a. stromal cell-derived factor 1, SDF-1), TGFB2, or PDGFB in our EPIC-ECM samples.

On a final note, our work also identified proteins present in both EPIC-ECM and EPIC-EVs, such as ANXA2, ANXA1, FN, HTRA1, CXCL12, EMILIN-1 and MYH9. Relevant to this finding, several studies have shown that EVs can influence cellular behavior as well as ECM modulation and reorganization (110–112). Importantly, matricellular proteins are thought to be internalized by endocytosis and re-secreted as EVs (112). This may hint at the existence of an epicardial ECM-EV interplay in the heart, which may ultimately affect autocrine and paracrine EV functions.

Altogether, the results of our study provide novel experimental evidence for the essential contribution of epicardium and epicardial-derived cells to the cardiac extracellular environment. The data presented here has clinical translational potential at different levels. Our data may contribute to using EVs as *bona fide* diagnostic or prognostic biomarkers for cardiac health (37, 74, 113–115). Translational studies involving EVs used as vectors have grown exponentially during the last decade, giving rise to the promise of EV-based therapies for the repair/regeneration of the diseased heart (reviewed in (33). Finally, the use of biocompatible matrices, alone or in combination with cells, and growth factors remains a plausible strategy for the repair of damaged tissues (116). Due to its characteristic protein content and its similarities with the embryonic cardiac ECM, the epicardial-secreted ECM described in our study could be of interest for developing biological scaffolds for reparative therapies.

