## Supplementary material for "Revealing the Complexity of the Epicardial Secretome: Characterization and Functional Assessment of Epicardium-Derived Extracellular Vesicles and Matrix": Figure Legends

**Figure 1 – Characterization of EPIC-derived EVs.** A) Representative TEM images of EVs-N (i), EVs-H5% (ii), and EVs-H1% (iii) samples. Scale bar: 50 nm; B) Particle concentration (number of particles/mL) of EVs-N (7.7 ± 2.3 x $10^{11}$ particles/mL), EVs-H5% (3.5 ± 0.5 x $10^{11}$ particles/mL) and EVs-H1% (2.8 ± 0.4 x $10^{11}$ particles/mL); C) Modal sizes of EVs-N (142.2 ± 25.5 nm), EVs-H5% (136.5 ± 13.1 nm) and EVs-H1% (103.8 ± 10.5 nm); D) Distribution of particle population in three different size ranges from EVs-N (35 – 200 nm: 73.8 ± 2.2%; 200 – 400 nm: 19.6 ± 1.9%; > 400 nm: 2.5 ± 0.6%), EVs-H5% (35 – 200 nm: 80.1 ± 3.3%; 200 – 400 nm: 17.8 ± 1.9%; > 400 nm: 2.1 ± 0.4%) and EVs-H1% (35 – 200 nm: 86.4 ± 1.7%; 200 – 400 nm: 12.3 ± 1.6%; > 400 nm: 1.3 ± 0.3%); E) Representative images of western blot against ALIX (95 kDa) and TSG101 (45 kDa) accumulation in protein lysates from total mouse embryos at E11.5 (control), EVs-N, EVs-H5%, EVs-H1%. Data are presented as mean ± SEM, n=3. One-way ANOVA test for particle concentration and modal size. Two-way ANOVA test for particle size distribution, *p < 0.05.

**Figure 2 – Proteomic analysis of epicardial-derived EVs isolated from EPIC cultured in normoxic or hypoxic conditions.** A) PCA representation shows identified proteins from EVs-N ("Normoxia_0.norm", "Normoxia_1.norm" and "Normoxia_2.norm"), EVs-H5%, ("Hypoxia_5_3.norm", "Hypoxia_5_4.norm" and "Hypoxia_5_5.norm"), and EVs-H1% ("Hypoxia_1_6.norm", "Hypoxia_1_7.norm" and "Hypoxia_1_8.norm"); B) Heatmap representation of identified proteins that significantly differ between EVs-H5% or EVs-H1% and EVs-N (p-value < 0.05; n=3). Scale represents standardized $\log_2$ of the protein abundancies with a mean of zero and standard deviation of one; C); Ten most representative GO annotations of biological process from significantly up-regulated proteins in EVs-H1% as compared to EVs-N. The x-axis represents the protein ratio, dot size corresponds to the number of proteins associated with the functional category, and dot colour corresponds to the adjust p value < 0.05D) Graphical representation of the fold change increase of glycolysis-related protein abundancy in EVs-H5% and EVs-H1% in respect to EVs-N. The inner circle corresponds to a fold change (FC) of 1 and the outer circle to an FC of 2.

**Figure 3 – Autocrine and paracrine effect of the cargo of epicardial-derived EVs.** A) Representative image of EPICs after incubation with EVs stained with DiI (EVs-DiI). (i) Bright-field image of EPIC nucleus stained with DAPI containing EVs-DiI (red); frames of time-lapse of EPIC stained with Lysotracker (acidic organelles in green) after the incubation with EVs-DiI at time 0 seconds (ii), 72 seconds(iii) and 154 seconds (iv); B) Representative images of HUVEC incubated with PKH26-stained EVs. (i) and (iii) show bright field images of HUVEC nucleus stained with DAPI containing PKH26-stained EVs (red); (ii) and (iv) show HUVEC containing PKH26-stained EVs, acidic organelles stained with Lysotracker (green) and nuclei stained with DAPI. Scale bars: 10 µm. C) Percentage of $EdU^+$ EPICs incubated for 24 hours at basal condition (grey bar). 20 µg/mL (yellow bars) or 50 µg/mL (blue bars) of EVs-N, EVs-H5% or EVs-H1% (*p < 0.05, **p < 0.01, ****p < 0.0001 against basal condition); D) Percentage of $EdU^+$ HUVECs incubated for 24 hours at basal condition (black bar). 20 µg/mL (yellow bars) or 50 µg/mL (blue bars) of EVs-N, EVs-H5% or EVs-H1% ; E) PER profiles of EPICs or (F) HUVECs incubated with 50 µg/mL of EVs-N, EVs-H5% or EVs-H1% or at basal condition (n=3); G) Glycolysis stress test parameters (basal acidification, glycolysis, glycolytic capacity, and glycolytic reserve) were calculated for EPICs and (H) HUVECs incubated with 50 µg/mL of EVs-N, EVs-H5% or EVs-H1% or at basal condition (n=3).

**Figure 4 – EPIC-derived ECM characterization and function assessment**. A) representative SEM images of EPIC and EPIC-derived ECM at 14,000X and 10,000X magnification, respectively; B) Venn diagram of extracellular proteins identified in EPIC IM, EPIC SM, and E17.5 hearts; C) Biological processes (gene ontology analysis) of extracellular proteins in common between EPIC IM, EPIC SM, and E17.5 hearts. The most highly significant categories in descending order in categories of gene ratio, are defined as the proportion of significant genes that are found in the functional category. The x-axis represents the gene ratio and the dot size the number of genes associated with the functional category and the dot color corresponds to the adjust p value<0.05.

**Figure 5 – EPIC-derived ECM function assessment.** HUVECs were incubated with 1% gelatine, 1% gelatine + 100 µg/mL of EPIC IM, 1% gelatine + 100 µg/mL of EPIC SM, 1% gelatine + 100 µg/mL Matrigel, 1% gelatine + 100 µg/mL Matrigel + 100 µg/mL of EPIC IM, or 1% gelatine + 100 µg/mL of EPIC IM + 100 µg/mL of EPIC SM for three and five days. A) Percentage of EdU positive HUVECs after incubation for three days; B) Percentage of EdU positive HUVECs after incubation for five days; *$p < 0.05$, **$p < 0.01$, ***$p < 0.001$, ****$p < 0.0001$.
