## Supplementary material for "Revealing the Complexity of the Epicardial Secretome: Characterization and Functional Assessment of Epicardium-Derived Extracellular Vesicles and Matrix": Suppl Material

These Supplemental Data contain:

1- Supplemental Material
2- Supplemental Figures

Supplementary material

**Human Umbilical Vascular Endothelial Cells culture**

A density of 6.25-7x$10^3$ human umbilical vascular endothelial cells (HUVECs)/$cm^2$ (passage 6 to 8, kindly donated by Dr. Beatriz Martinez Poveda) were cultured in 1% gelatine-coated flasks and incubated at 37 °C and 5% $CO_2$ until they reached 80% confluency. HUVEC culture media was composed by Endothelial Cell Growth Basal Medium-2 (EBM-2, BulletKit™, Lonza) supplemented with 2% FBS, 0.1% heparin, 0.1% VEGF, 0.1% Ascorbic acid, 0.1% Gentamicin sulfate-Amphotericin (GA-1000), 0.1% R3-insulin-like growth factor-1 (R3-IGF-1), 0.1% recombinant human Epidermal Growth Factor (rhEGF) and 0.04% hydrocortisone, 0.4% recombinant human Fibroblast Growth Factor-β (rhFGF-β) (all from Lonza), 100 U/mL penicillin (Sigma), 100 µg/mL streptomycin (Sigma).

**Visualization of fluorescent-labeling extracellular vesicles**

EPIC-derived EVs were probed with two different fluorophores. CellTracker™ CM-DiI (Invitrogen) was added together with f-PBS during EPIC-derived EVs isolation and ultracentrifugation at 100,000 g for 70 min (4 °C). The supernatant was discarded, and DiI-probed EVs were resuspended in f-PBS and stored at -80 °C. Alternatively, 3 µg of EVs were labeled with 1 uL of PKH26 (Sigma) following manufacturer's instructions. Then, 95 µL of Diluent C was added to both samples and incubated for 5 min at RT, followed by the addition of 100 µL f-PBS (5 min at RT). Finally, the samples were centrifuged at 10,000 g for 60 minutes (25 °C), and the supernatant was collected into a new tube and centrifuged at 16,000 g for an additional period of 30 minutes. All samples were stored at -80°C. Both CellTracker™ CM-DiI and PKH26 were diluted in f-PBS without EVs as negative controls. For visualization of the vesicles, 20 uL of stained EVs and negative controls were imaged using Eclipse Ti TIRF Microscope (Nikon). Images were processed using ImageJ 1.53i software.

**Internalization assays**

To study EPIC-derived EVs internalization in EPIC, 30 µL from either a 1/100 diluted DiI-probed EVs solution, PBS containing DiI, or PBS alone, were diluted in an EV-depleted medium and added to a 70% confluence flask. To assess EPIC-derived EVs internalization in HUVEC, 40 µg of PKH26-dyed EVs, PBS containing PKH26, or PBS alone were diluted in EBM-2 and added to HUVECs at 70% of confluency. EPIC and HUVEC were then incubated at 37 °C and 5% $CO_2$ for 4 hours. Hoechst 33342 dye (1,5 µg/mL) (Thermo Fisher) and LysoTracker® Green DND-26 (50 nM) (Thermo Fisher) were added to stain nuclei and lysosomes, respectively. Samples were then imaged using Eclipse Ti TIRF Microscope (Nikon) and acquired data were processed using ImageJ 1.53i software.

**Proliferation assay**

For EPIC and HUVEC cultures incubated with EPIC-derived EVs or plated on EPIC IM- and EPIC SM-coated plastic dishes, cells were incubated for 15 min at 37 °C with 10 µM of 5-Ethynyl-2′-deoxyuridine (EdU), fixed with 4% paraformaldehyde (PFA, Sigma), permeabilized with 0.5% of Triton X-100, and washed in DPBS. Then, cells were incubated with the Click reaction mixture for 15 min at RT. To quantify final cell numbers, plates were incubated with 4',6-diamidino-2-phenylindole (DAPI, 1:4,000 in PBS (15 min, RT). Cell proliferative rates were analyzed using the Operetta High Content Screening system (Perkin Elmer) and Harmony 4.8 software. DAPI and Alexa labeled nuclei were counted and the percentage of EdU-positive cells per well was calculated using the following formula:

$$(Number\ of\ EdU\ positive\ nucleus)/(Number\ of\ DAPI - labelled\ nucleus)\ X\ 100$$

**Glycolysis stress assay induced by EPIC-derived extracellular vesicles**

EPIC or HUVEC were plated in Seahorse XF24 cell culture plates (Agilent) (28,000 cells/well). For HUVEC plating, wells were previously coated with 1% gelatine. After 48 hours, cells were washed in warm PBS and incubated overnight in 0% FBS medium for EPIC or 1% FBS medium for HUVEC. The following day, starving media was replaced with 20 or 50 µg/mL of EVs-N, EVs-H5%, or EVs-H1, and cells were incubated for 12 hours at 37 °C and 5% $CO_2$. The following day, cells were incubated for 1 hour at 37 °C in a non-$CO_2$ incubator in a pyruvate-free glycolytic assay medium composed of XF base media (Agilent) with 2 mM glutamine (Sigma), pH 7.4. Then, cells were sequentially incubated with 10 mM D-glucose, 1 µM Oligomycin A, 50 mM 2-DG, and 2 mM L-glutamine, following the manufacturer's instructions. After three baseline measurements, glycolytic parameters were calculated with three cycles of measurements after each injection. At the end of the assay, supernatants were removed, and the cells lysed with RIPA for protein quantification (see details below). Values were normalized to protein quantity measured by BCA quantification assay and data was analyzed with Wave 2.6.1 software (Agilent) and GraphPad 8.0.2.

**Scanning Electron Microscopy**

A total of 7-8 $x10^5$ EPIC/$cm^2$ were seeded onto 24 mm coverslips (Thermo Fisher) in 100-mm dishes. EPIC and EPIC IM were fixed in 2% (v/v) glutaraldehyde (Sigma) for 1 hour at RT. After fixation, cell and ECM samples were dehydrated using an increasing concentration of ethanol (from 30% up to 100%) and deposited in an aluminum drum with a conductive double-sided adhesive tape. Samples were sputter-coated with a thin layer of gold (300 Å) in a Sputtering QOURUM Q 150R (QOURUM). Samples were observed using JEOL JSM6490LV SEM (JEOL) operating at 15 kV and a working distance of 10 mm.

**Immunocytochemistry**

Coverslips containing EPIC and EPIC IM were fixed with 4% PFA (for 20 and 10 minutes, respectively), whereas EPIC SM pellets were fixed in 4% PFA for 30 minutes after being washed in PBS and centrifugated. Fixed ECM fractions were blocked with SBT (10% horse serum, 1.5% BSA, and 0.5% Triton X-100 in Tween-PBS) for 1 hour and incubated overnight at 4 °C with the primary antibodies, either 1:100 rabbit polyclonal anti-fibronectin (Sigma) or 1:200 monoclonal anti-Laminin subunit α1 (Sigma). SBT without any primary antibodies was used as negative control. Samples were then incubated with DAPI (1:2,000 in PBS) and anti-rabbit AF647 (1:200 in PBS; Jackson Immuno Research) for 1 hour at RT. For F-actin staining, EPIC and EPIC ECM were incubated with 0.6 µM of Phalloidin Atto 488 (Thermo Fisher) and DAPI (1:2,500 in PBS) for 20 minutes in a rocking shaker. Coverslips were mounted on microscope glass slides with PBS: Glycerol (1:1). All samples were analyzed using Leica SP5 HyD Confocal Microscope and images were processed using ImageJ 1.53i software.

**Protein quantification**

EPIC protein lysates were prepared using RIPA buffer (5 mM of Trizma, 15 mM NaCl, 0,1% Triton X-100, 0,1 mM EDTA, and 0,6 mM of sodium deoxycholate in distilled water) containing 1% of protease inhibitor cocktail (SIGMA). The cell lysate was then incubated on ice for 5 minutes and centrifuged at 12,000 g, 4 °C for 10 min and the supernatant was collected. For the quantification of the EVs samples, 10 µL EVs were lysed in 6 µL of lysis buffer composed of

1% Triton-X100 and 0,1% sodium dodecyl sulfate (SDS) (SIGMA) in distilled water for 30 min on ice. For ECM protein quantification, EPIC IM was thoroughly detached in Laemmli buffer (4% SDS, 20% glycerol, 10% 2-mercaptoethanol, 0.004% bromophenol blue and 0.125 M Tris HCl) using cell scrapers. EPIC SM was boiled for 15 min, vortexed every 2 min, sonicated in an ultrasonic bath for 5 min, and centrifuged at 14,000 g for 5 min. Both fractions were then homogenized in SDS-PAGE buffer at 95 ºC. All samples were stored at -80 ºC. EPIC-derived EVs and ECM protein quantification were performed using the Pierce™ BCA Protein Assay Kit (Thermo Fisher) following the manufacturer's instructions for cell lysate (adapted to 2h, 60 °C).

**Western Blot**

Either a total of 16 µg (cell lysates and EVs samples) or 10 µg (ECM samples) of protein were separated in 8-12% Bis/Tris polyacrylamide gels. Proteins were transferred onto nitrocellulose membranes (Protan®, 0.45 µm). To confirm protein transference and serve as protein loading control, a Ponceau staining (5% (m/v) Ponceau Red in 0.1% (v/v) acetic acid) was performed. Then, membranes were blocked using 5% (m/v) non-fat dry milk diluted in TBS-T (10 mM Tris-HCl, 150 mM NaCl, 0,005% Tween-20) for 1 h at RT, and incubated overnight with agitation at 4°C (primary antibodies diluted in blocking buffer). EPIC lysates and EPIC-derived EV proteins were incubated with mouse anti-TSG101 (1:100; Santa Cruz Biotechnology) or anti-ALIX (1:100, Santa Cruz Biotechnology). For EPIC-ECM analysis, proteins were incubated with mouse anti-Laminin subunit α1 (1:1,000; SIGMA) or rabbit anti-fibronectin (1:1,000; SIGMA). After three washes with TBS-T, HRP-conjugated anti-mouse IgG (1:4,000; SIGMA) and anti-rabbit IgG antibodies (1:10,000; SIGMA) diluted in TBS-T were added to membranes for 1 h at RT. Membranes were revealed using the SuperSignal® West Pico PLUS Chemiluminescent Substrate reagent (ThermoFisher) according to the manufacturer's instructions, imaged using the ChemiDoc XRS+ imaging system (BIO-RAD) and analyzed with Image Lab software (BIO-RAD).

**Proteome analysis of EPIC-derived extracellular vesicles by multiplexing proteomics**

Tandem Mass Tags (TMT) were employed to allow the analysis of multiple samples in the same assay, diminishing the variability between technical replicates. Proteins extracted from EVs (EVs-N, EVs-H5% and EVs-H1%, n=3 each condition) were diluted to 1ug/uL and incubated with 200 mM tris (2-carboxyethyl)phosphine (TCEP) for 1 h at 55 °C. Then, samples were incubated for 30 min with tetraethylammonium bromide (TEAB) containing 60 mM of iodoacetamide, precipitated in acetone at −20 °C for 4 h, and centrifugated at 8,000 g for 10 min at 4 °C. Pellets were resuspended in 50 mM TEAB (pH 8.5) and incubated in trypsin overnight at 37 °C. Protein samples (500 ng) were tagged using a TMT10plex Isobaric Mass Tagging Kit (Thermo Fisher Scientific) according to the manufacturer's instructions. The chemically tagged samples were combined into one tube and dried using SpeedVac. Purified samples were resuspended with 0.1% of formic acid (FA) and analyzed by LC-MS/MS.

**Descriptive composition of EPIC-derived ECM by label-free proteomics**

Qualitative analysis of ECM composition was performed by label-free bottom-up mass spectrometry. EPIC IM (n=3), EPIC SM (n=3), and decellularized E17.5 hearts (n=6) fractions were solubilized in SDS-PAGE buffer. EPIC-derived ECM, embryonic hearts, and Matrigel® (Corning) (n=3) were denatured at 95 C for 15 min, vortexed intermittently, sonicated for 5 min, and centrifuged at 14, 000 g for 5 min. To eliminate incompatible components of LC-MS, the protein solution was entrapped in a polyacrylamide gel matrix before reduction with dithiothreitol and cysteine carbamidomethylating with iodoacetamide. Gels were cut into 1-2 mm cubes and treated with 50% acetonitrile (ACN)/25 mM ammonium bicarbonate. Samples were dehydrated, resuspended with ACN, and reduced with 10 mM DTT in 50 mM ammonium bicarbonate for 30 min at 56 °C. Cysteine residues were carbamidomethylated with 55 mM iodoacetamide in 50 mM of ammonium bicarbonate for 20 min at RT. Gel fractions were dehydrated, and proteins were digested by rehydrating gel fractions in 10 ng/µL trypsin (Promega) overnight at 30 ºC. Peptides were extracted with FA/ACN (0.1%/80%) for 30 min at RT. Samples were dried using SpeedVac, dissolved in 0.1% FA, treated with ultrasound for 3 min, centrifuged at 13, 000 g for 5 min, and finally purified and concentrated using C18 ZipTip (Merck) according to manufacturer's instructions. All samples were analyzed by LC-MS/MS.

**Liquid chromatography-tandem mass spectrometry (LC-MS/MS)**
For LC-MS/MS analysis, purified samples were injected into an Easy nLC 1200 UHPLC coupled to Q Exactive HF-X Hybrid Quadrupole-Orbitrap mass spectrometer (ThermoFisher Scientific). Mobile phases of HPLC consisted of 0.1% FA (in water) and FA/acetonitrile (ACN) (0.1%/80%). Using a thermostatic, automatic injector, 100 ng of the peptide sample was loaded into a pre-column Acclaim PepMap 100, 75 µm x 2 cm, C18, 3 µm, 100 A (ThermoFisher Scientific) at a flow of 20 µL/min and separated in a 50 cm analytical column (PepMap RSLC C18, 2 µm, 100 A, 75 µm x 50 cm; ThermoFisher Scientific).

ECM-derived peptides were eluted from the analytical column with a 120-min gradient from 5% to 20% FA/ACN (0.1% FA/80% ACN), followed by a 5-min gradient from 20% to 32%, and finally to a 95% for 10 min before rebalancing with 5% FA/ACN (0.1%/80%). MS scans were performed in the 375-1,600 m/z range at a resolution of 120,000 m/z. Using a data-dependent acquisition mode, the 15 strongest precursor ions were isolated from all precursor ions with a charge of +2 to +5 within a 1.2 m/z window and fragmented to obtain the corresponding $MS^2$ spectra. Fragmentation ions were generated in a high energy collisional dissociation cell (HCD) with a fixed first mass at 110 m/z and detected on an orbitrap mass analyzer at a resolution of 30,000. The dynamic exclusion for the selected ions was 30 sec. The maximum allowed ion accumulation time in MS and $MS^2$ mode was 50 msec and 70 msec, respectively. Automatic gain control was used to prevent overfilling of the ion trap and was set to $3x10^6$ ions and $2x10^5$ ions for a full MS and $MS^2$ scan, respectively.

EV-derived peptides tagged with TMT were eluted with a gradient of 2 to 20% of FA/ACN (0.1%/80%) for 4 h. This step was followed by elution with a 20 to 35% gradient of FA/ACN (0.1%/80%) for 30 min and finally a gradient of 95% of FA/ACN (0.1%/80%) for 15 min before rebalancing the column with 2% of FA/ACN (0.1%/80%). For both ECM and EV peptides, the elution was performed at a constant flow of 300 nL/min. The resolution of MS survey scans was set to 120,000 m/z, while for MS/MS the resolution was set to 30,000. The ion pulverization voltage was set to 2.2 kV with an m/z window from 350 to 1,500, an isolation window of 0.7 m/z, and dynamic exclusion of 20 sec. Software versions used for data acquisition and manipulation were Tune 2.9 and Xcalibur 4.1.31.9.

**Bioinformatic analysis**
Searches of the $MS/MS^2$ spectra were performed against the SwissProt *Mus musculus* version 2017.10.25 protein database (25,097 sequences). The raw data acquired was analyzed on the Proteome Discoverer 2.2 platform (Thermo Fisher Scientific) with the Sequest HT search engine set for mass tolerances of 10 ppm and 0.02 Da for the precursor ions and fragment ions, respectively. Two lost tryptic cleft sites were allowed. Oxidation of methionine and N-terminal acetylation were established as variable modifications, while carbamidomethylation of cysteine residues was established as a fixed modification. Exploratory protein identification

was pursued by identifying protein presence or absence in two or more biological samples with three replicates for EPIC IM, EPIC SM, and Matrigel® and six replicates for E17.5 decellularized hearts. Proteins were considered part of an experimental condition if they were present in at least two replicates. The false discovery rate (FDR) for consecutive protein and peptide assignments was determined using the Percolator software package, based on a target-decoy approach that uses an inverted protein database as a decoy, imposing a strict limit of 1% FDR. The results were filtered to accept only those proteins with at least two peptide sequences.

For TMT data analysis, *MaxQuant* (Version 1.5.4.1) software was used with RAW files and the SwissProt mouse protein database (release 2019_11) supplemented with common contaminants (Cox and Mann, 2008; The Uniprot Consortium, 2023). Parameters in Andromeda search engine for EVs tagged with TMT include 20 ppm peptide precursor mass tolerance and 0.5 Da fragments mass tolerance (Cox *et al.*, 2011). The search space also included two sites of tryptic excision. Oxidation of methionine and N-terminal acetylation were allowed as variable modifications and carbamidomethylation of cysteine was set as fixed modification. The rate of false positives (FDR) was determined using a target-decoy strategy and was kept at 1% at the peptide and protein level. Protein identifications related to sample contaminants, DECOY peptides, or without quantification were eliminated. Proteins that were not identified in, at least, two samples of each experimental condition, would not be considered for the statistical analysis. Protein abundances were transformed using Log2 and normalized by quantile normalization.

**Comentado [AR1]:** Para Eli.
Uno de los revisores e ha quejado de esto:
"Reviewer 1: "the number of entries in the SwissProt mouse database (release 2019_11) were actually searched".
¿Qué quiere decir?

For ECM-derived peptide analysis, relative abundances of sample replicates were prepared by calculating the abundance of a protein relative to total abundance per condition. In detail, and to obtain a percentage for the relative abundance of each protein, relative abundances were calculated per protein as follows:

$$(Average\ of\ protein\ abundance)/(Total\ proteins\ abundance)\ \ x\ 100$$

Total abundance was calculated from the sum of abundances per replicate. Then, the average of replicate total abundance was calculated to obtain the total abundance per condition. A list of proteins and respective abundances were then filtered in the "cellular component" tab for the "extracellular" term. For ECM analysis, only extracellular proteins were considered.

All raw data are available under request to the corresponding authors.

Supplementary Figures

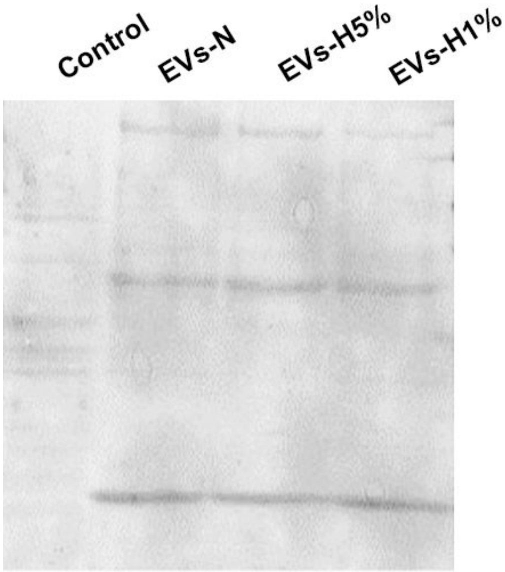


**Figure S1 –** Ponceau staining. Western blot analysis of a total protein extract from a pool of E11.5 embryo tissues (control), EVs-N, EVs-H5% and EVs-H1%. Cell lysates were tested against TSG101 and ALIX.

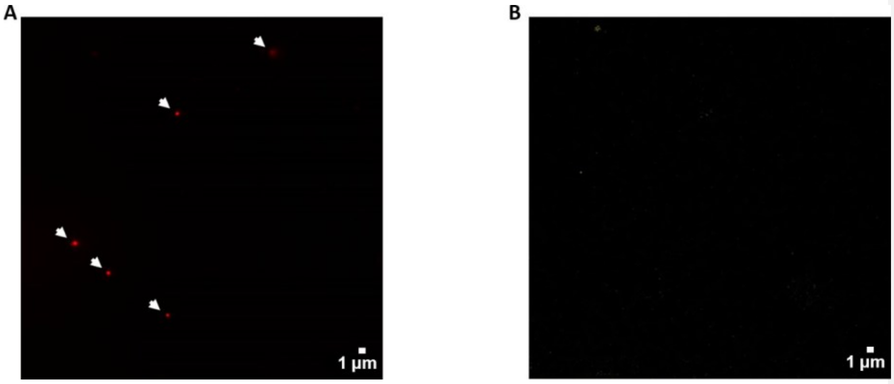


**Figure S2 –** Stainning of EPIC-derived EVs. A) EPIC-EVs stainned with 0.8 µM DiI, washed with f-PBS and shown after ultracentrifugation; B) Ultracentrifuged DiI solution in PBS (0.8 µM), used as negative control.

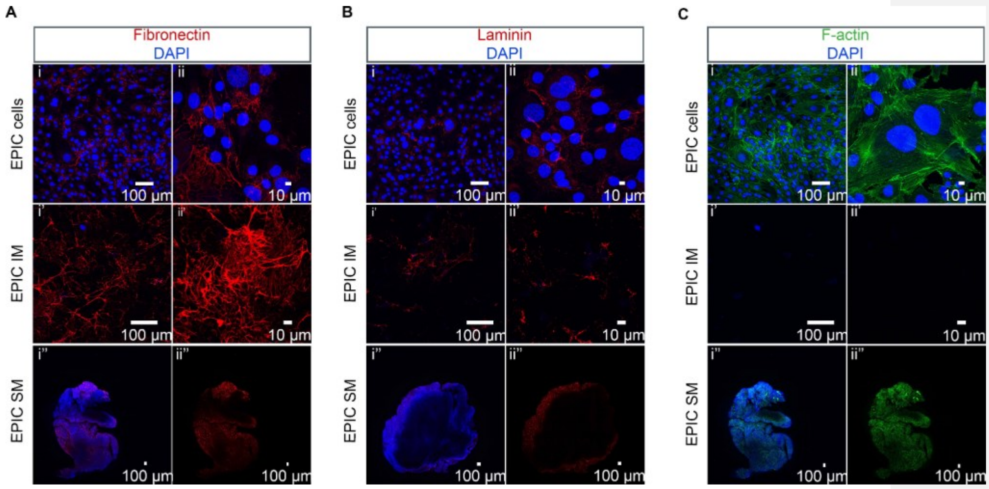


**Figure S3 -** Characterization of EPIC, EPIC IM and EPIC SM. Immunohistochemical analysis of ECM proteins such as fibronectin (A) and laminin subunit α1 (B). Cytoskeletal protein F-actin detection (C). In all conditions, nuclear content was stained with DAPI. EPIC cells, EPIC IM fraction and EPIC SM fraction are represented in images i and ii, i' and ii', and i'' and ii'', respectively.

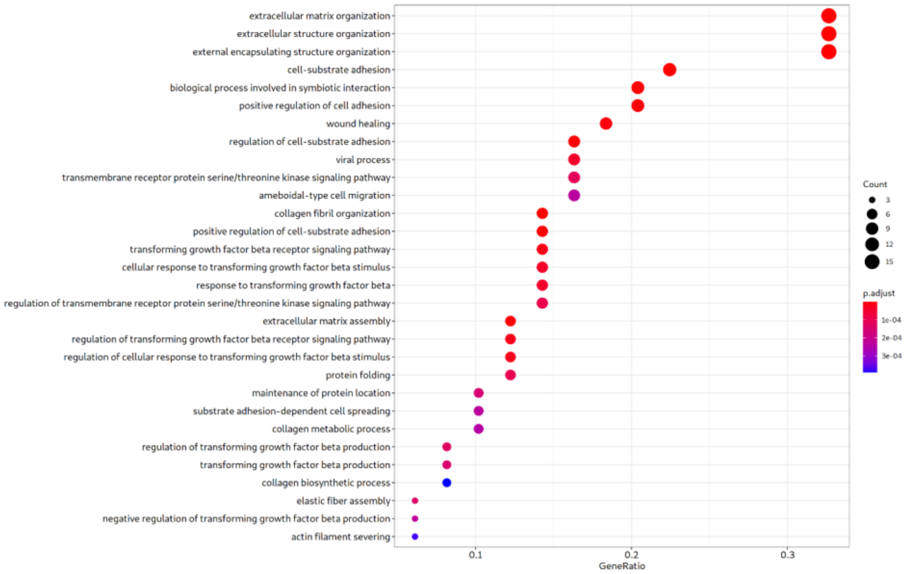


**Figure S4 -** Biological processes (gene ontology analysis) of extracellular proteins present in both EPIC IM and EPIC SM represented in a dot plot graph. The x-axis represents the protein ratio and the dot size the number of genes associated with the functional category; the colour indicates the p.adjust value.

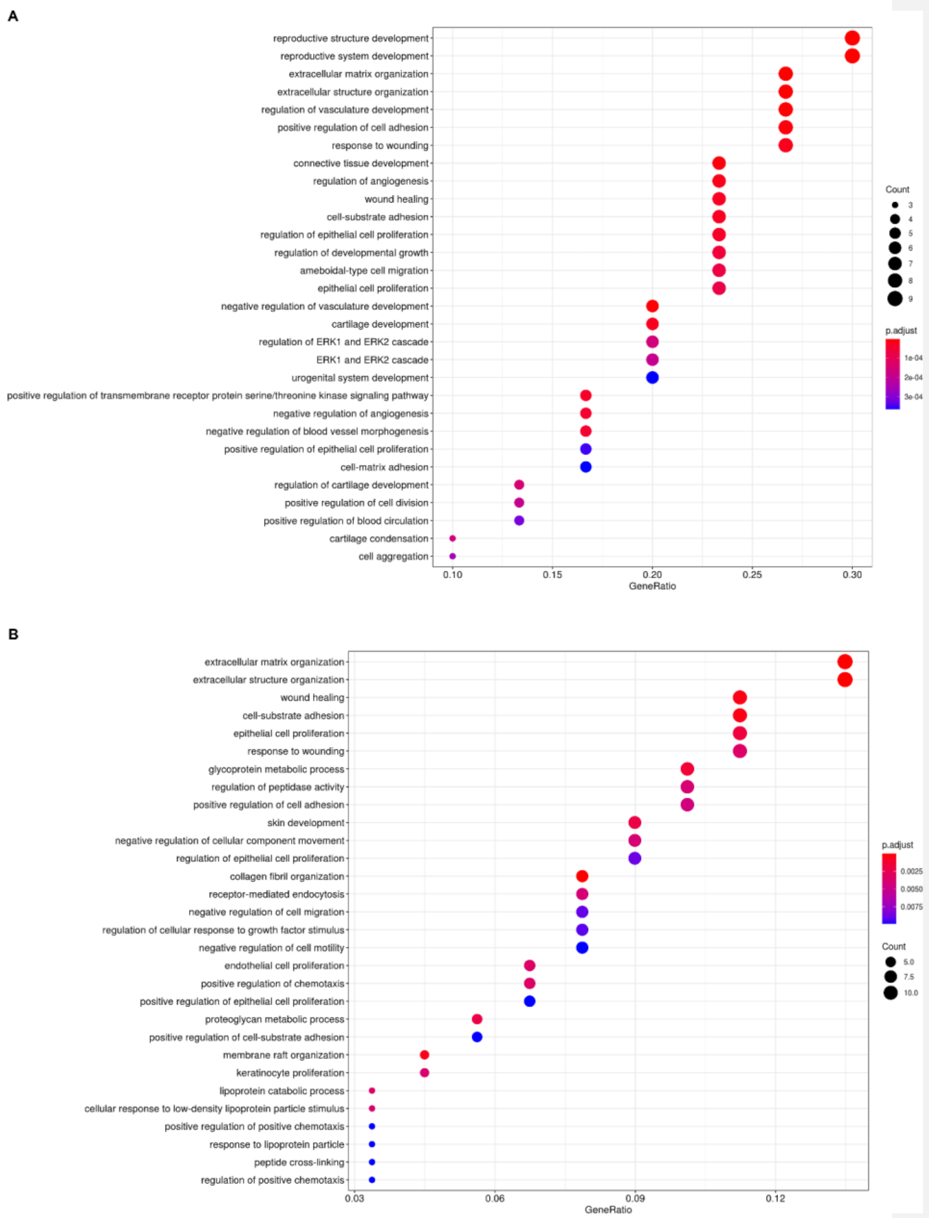


**Figure S5 -** Biological processes (gene ontology analysis) of A) EPIC IM unique extracellular proteins and B) EPIC SM unique extracellular proteins represented in a dot plot graph. The x-axis represents the protein ratio and the dot size the number of genes associated with the functional category; the colour indicates the p.adjust value.

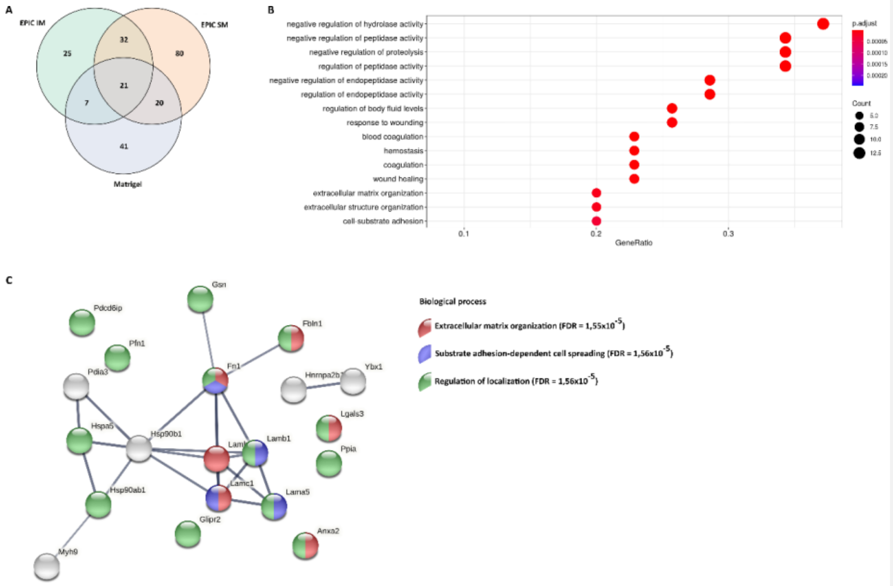


**Figure S6** - Qualitative comparison of identified proteins in a label-free proteomic analysis of EPIC IM, EPIC SM and Matrigel samples. A) Venn diagram representation of extracellular proteins identified in EPIC IM (85), EPIC SM (153) and Matrigel (89). EPIC IM shares 7 extracellular proteins with Matrigel, while EPIC SM shares 20 extracellular proteins with Matrigel. Twenty-one proteins are shared between all samples; B) Biological processes (gene ontology analysis) of extracellular proteins from Matrigel represented in a dot plot graph. The x-axis represents the protein ratio and the dot size the number of genes associated with the functional category and the colour corresponds to the p.adjust value. C) STRING analysis of extracellular proteins shared between EPIC IM, EPIC SM and Matrigel, and the three biological process terms that show the lowest FDR values.

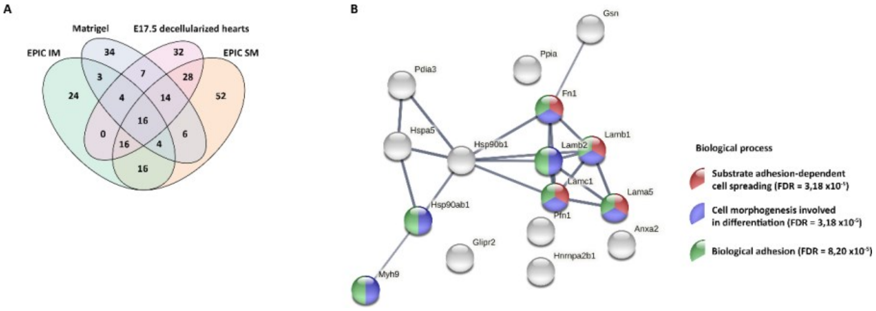


**Figure S7 -** Qualitative comparison of identified proteins in label-free proteomics from EPIC IM, EPIC SM, E17.5 decellularized hearts and Matrigel samples. A) Venn diagram representation of extracellular proteins in EPIC IM, EPIC SM E17.5 decellularized hearts and Matrigel. EPIC IM, EPIC SM, E17.5 hearts and Matrigel contain 25, 52, 32 and 34 unique proteins, respectively. Seventeen proteins are found to be common to all samples; B) STRING analysis of proteins shared between EPIC IM, EPIC SM, E17.5 hearts and Matrigel, and the three biological process terms that show the most significant FDR values.

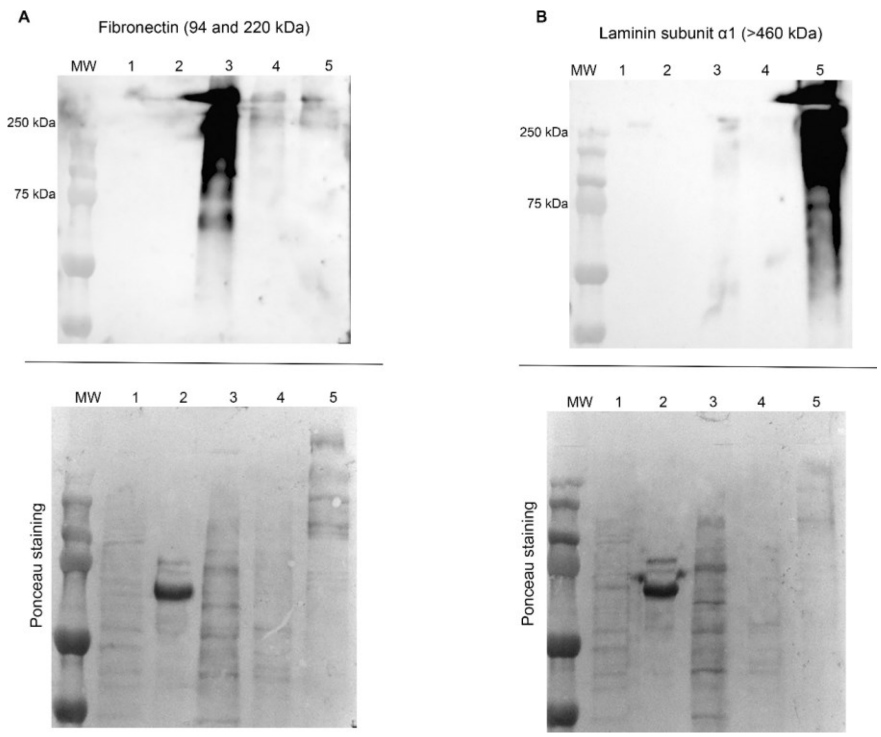


**Figure S8 -** Western Blot analysis of EPIC (lane 1), EPIC IM (lane 2), EPIC SM (lane 3), E17.5 heart cell lysates lysate (lane 4) and Matrigel (lane 5). Samples have been probed against fibronectin (A) and laminin subunit α1 (B) and the corresponding Ponceau staining is shown.
