## Supplementary material for "Revealing the Complexity of the Epicardial Secretome: Characterization and Functional Assessment of Epicardium-Derived Extracellular Vesicles and Matrix": Suppl Tables

### Supplementary tables

**Table S1 -** Extracellular proteins identified in both EPIC IM and EPIC SM fractions.

| UniProt ID | Protein name | UniProt ID | Protein name |
|---|---|---|---|
| O08573 | LGALS9 | P39876 | TIMP3 |
| O88569 | HNRNPA2B1 | P48759 | PTX3 |
| P01029 | C4B | P55065 | PLTP |
| P01887 | B2M | P62960 | YBX1 |
| P01902 | H2-K1 | P62962 | PFN1 |
| P02468 | LAMC1 | P82198 | TGFBI |
| P02469 | LAMB1 | P97792-1 | Isoform 1 of CXADR |
| P06745 | GPI1 | Q01149 | COL1A2 |
| P07356 | ANXA2 | Q05793 | HSPG2 |
| P08113 | HSP90B1 | Q08879 | FBLN1 |
| P10107 | ANXA1 | Q61001 | LAMA5 |
| P10649 | GSTM1 | Q61292 | LAMB2 |
| P11087-1 | Isoform 1 of COL1A1 | Q61398 | PCOLCE |
| P11152 | LPL | Q61703 | ITIH2 |
| P11276 | FN1 | Q62351 | TFRC |
| P11499 | HSP90AB1 | Q8CG19-1 | Isoform large of LTBP1 |
| P13020-1 | Isoform 1 of GSN | Q8R2G6 | CCDC80 |
| P16110 | LGALS3 | Q8VDD5 | MYH9 |
| P17742 | PPIA | Q924C6 | LOXL4 |
| P18760 | CFL1 | Q99K41 | EMILIN1 |
| P19788 | MGP | Q9CYL5 | GLIPR2 |
| P20029 | HSPA5 | Q9D1D6 | CTHRC1 |
| P21956-1 | Isoform 1 of MFGE8 | Q9D6X6 | PRSS23 |
| P27773 | PDIA3 | Q9R118 | HTRA1 |
| P28301 | LOX | Q9WU78 | PDCD6IP |
| P35441 | THBS1 | Q9WVJ9 | EFEMP2 |
| P37889 | FIBL-2 | | |

**Table S2 –** Extracellular proteins identified in EPIC IM or EPIC SM fractions only.

| Unique proteins from EPIC SM | | | | Unique proteins from EPIC IM | |
|---|---|---|---|---|---|
| **UniProt ID** | **Protein Name** | **UniProt ID** | **Protein Name** | **UniProt ID** | **Protein name** |
| A2ASQ1-1 | Isoform 1 of AGRN | Q00780 | COL8A1 | A2A5I3 | R3HDML |
| D3YXK1 | SAMD1 | Q02788 | COL6A2 | A6X935-1 | Isoform 1 of ITIH4 |
| E9Q414 | APOB | Q04857 | COL6A1 | O09118 | NTN1 |
| O08638-1 | Isoform 1 of MYH11 | Q07797 | LGALS3BP | P01027-1 | Isoform long of C3 |
| O08912 | GALNT1 | Q3U962 | COL5A2 | P07724 | ALB |
| O08992 | SDCBP | Q3V1T4 | P3H1 | P09535-2 | Isoform 2 of IGF2 |
| O35516 | NOTCH2 | Q5SS80-1 | Isoform 1 of DHRS13 | P10493 | NID1 |
| O35604 | NPC1 | Q60847 | COL12A1 | P11214 | PLAT |
| O35658 | C1QBP | Q60854 | SERPINB6A | P14069 | S100A6 |
| O35887 | CALU | Q61147 | CP | P18406 | CCN1 |
| O88207 | COL5A1 | Q61207 | PSAP | P19137 | LAMA1 |
| O88531 | PPT1 | Q61245-1 | Isoform long of COL11A1 | P24383 | WNT7A |
| O89051 | ITM2B | Q61592 | GAS6 | P27090 | TGFB2 |
| P02463 | COL4A1 | Q61699 | HSPH1 | P29268 | CCN2 |
| P07091 | S100A4 | Q62086 | PON2 | P31240 | PDGFB |
| P07214 | SPARC | Q62087 | PON3 | P40224-2 | Isoform beta of CXCL12 |
| P08121 | COL3A1 | Q62165 | DAG1 | P43028 | GDF6 |
| P08122 | COL4A2 | Q64437 | ADH7 | P63089 | PTN |
| P08207 | S100A10 | Q6GU68 | ISLR | P70275 | SEMA3E |
| P09528 | FTH1 | Q80ZW2 | THEM6 | P97298 | SERPINF1 |
| P10605 | CTSB | Q8C7K6 | PCYOX1L | P97857 | ADAMTS1 |
| P10810 | CD14 | Q8C7V8-1 | Isoform 1 of CCDC134 | Q3UQ28 | PXDN |
| P14211 | CALR | Q8CC88 | VWA8 | Q3USZ8 | DIPK2A |
| P16045 | LGALS1 | Q8CIE6 | COPA | Q62177 | SEMA3B |
| P18242 | CTSD | Q8K4Z3 | NAXE | Q66PY1-1 | Isoform 1 of SCUBE3 |
| P21460 | CST3 | Q8R422 | CD109 | Q6GQT1 | A2M |
| P21803-1 | Isoform Long of FGFR2 | Q8VDL4 | ADPGK | Q6R0H7 | GNAS |
| P23188 | FURIN | Q91VU0 | FAM3C | Q80T21 | ADAMTSL4 |
| P25446 | FAS | Q91ZA3 | PCCA | Q8K0E8 | FGB |
| P25785 | TIMP2 | Q920A5 | SCPEP1 | Q91V88-4 | Isoform 4 of NPNT |
| P28481-1 | Isoform 2 of COL2A1 | Q99104 | MYO5A | Q9JK53 | PRELP |
| P28653 | BGN | Q99JR5 | TINAGL1 | Q9Z0J7 | GDF15 |
| P28654 | DCN | Q99M71 | EPDR1 | | |
| P29391 | FTL1 | Q99MN1 | KARS | | |
| P30681 | HMGB2 | Q9CPT4 | MYDGF | | |
| P31230 | AIMP1 | Q9CXI5 | MANF | | |
| P32261 | SERPINC1 | Q9CYA0 | CRELD2 | | |
| P39061-1 | Isoform 2 of COL18A1 | Q9CYD3 | CRTAP | | |

| | | | |
|---|---|---|---|
| P40124 | CAP1 | Q9CZD3 | GARS |
| P40224-1 | Isoform alpha of CXCL12 | Q9D0L4-1 | Isoform 1 of ADCK1 |
| P40240 | CD9 | Q9DCZ4 | APOO |
| P41731 | CD63 | Q9EQ06 | HSD17B11 |
| P42703-1 | Isoform 1 of LIFR | Q9ET22 | DPP7 |
| P51655 | GPC4 | Q9JHR7 | IDE |
| P51859 | HDGF | Q9JKR6 | HYOU1 |
| P55302 | LRPAP1 | Q9QXP7 | C1QTNF1 |
| P63094 | GNAS | Q9QZF2 | GPC1 |
| P63158 | HMGB1 | Q9WTR5 | CDH13 |
| P81117 | NUCB2 | Q9Z204 | HNRNPC |
| P97352 | S100A13 | Q9Z2W0 | DNPEP |

**Table S3 –** Extracellular proteins common to DMEH, EPIC IM and EPIC SM extracts.

| **Proteins in common between EPIC IM and DMEH** | | **Proteins in common between EPIC SM and DMEH** | | **Proteins in common between EPIC IM, EPIC SM and DMEH** | |
|---|---|---|---|---|---|
| **UniProt ID** | **Gene Name** | **UniProt ID** | **Gene Name** | **UniProt ID** | **Gene Name** |
| P07724 | ALB | A2ASQ1-1 | Isoform 1 of AGRN | O08573 | LGALS9 |
| P10493 | NID1 | E9Q414 | APOB | O88569 | HNRNPA2B1 |
| Q3UQ28 | PXDN | O08638-1 | Isoform 1 of MYH11 | P02468 | LAMC1 |
| Q8K0E8 | FGB | O35604 | NPC1 | P02469 | LAMB1 |
| | | O35658 | C1QBP | P06745 | GPI1 |
| | | O35887 | CALU | P07356 | ANXA2 |
| | | P02463 | COL4A1 | P08113 | HSP90B1 |
| | | P08122 | COL4A2 | P10107 | ANXA1 |
| | | P14211 | CALR | P11087-1 | Isoform 1 of COL1A1 |
| | | P28653 | BGN | P11152 | LPL |
| | | P28654 | DCN | P11276 | FN1 |
| | | P29391 | FTL1 | P11499 | HSP90AB1 |
| | | P30681 | HMGB2 | P13020-1 | Isoform 1 OF GSN |
| | | P31230 | AIMP1 | P17742 | PPIA |
| | | P39061-1 | Isoform 2 of COL18A1 | P18760 | CFL1 |
| | | P40124 | CAP1 | P20029 | HSPA5 |
| | | P51655 | GPC4 | P21956-1 | Isoform 1 of MFGE8 |
| | | P63094 | GNAS | P27773 | PDIA3 |
| | | P63158 | HMGB1 | P28301 | LOX |
| | | Q02788 | COL6A2 | P35441 | THBS1 |
| | | Q04857 | COL6A1 | P37889 | FBLN2 |
| | | Q3V1T4 | P3H1 | P62962 | PFN1 |
| | | Q5SS80-1 | Isoform 1 of DHRS13 | P97792-1 | Isoform 1 of CXADR |
| | | Q60847 | COL12A1 | Q01149 | COL1A2 |
| | | Q61207 | PSAP | Q05793 | HSPG2 |
| | | Q62087 | PON3 | Q61001 | LAMA5 |
| | | Q62165 | DAG1 | Q61292 | LAMB2 |
| | | Q6GU68 | ISLR | Q61703 | ITIH2 |

| | | | |
|---|---|---|---|
| Q8CC88 | VWA8 | Q62351 | TFRC |
| Q8CIE6 | COPA | Q8CG19-1 | Isoform long of LTBP1 |
| Q8K4Z3 | NAXE | Q8VDD5 | MYH9 |
| Q8VDL4 | ADPGK | Q99K41 | EMILIN1 |
| Q91ZA3 | PCCA | Q9CYL5 | GLIPR2 |
| Q99MN1 | KARS | | |
| Q9CYD3 | CRTAP | | |
| Q9CZD3 | GARS | | |
| Q9D0L4-1 | Isoform 1 of ADCK1 | | |
| Q9DCZ4 | APOO | | |
| Q9JKR6 | HYOU1 | | |
| Q9QZF2 | GPC1 | | |
| Q9WTR5 | CDH13 | | |
| Q9Z2W0 | DNPEP | | |

**Table S4 -** Extracellular proteins shared by EPIC ECM fractions, Matrigel and decellularized embryonic murine heart (DMEH) samples.

| UniProt ID | Protein name |
|---|---|
| O88569 | HNRNPA2B1 |
| P02468 | LAMC1 |
| P02469 | LAMB1 |
| P07356 | ANXA2 |
| P08113 | HSP90B1 |
| P11276 | FN1 |
| P11499 | HSP90AB1 |
| P13020-1 | Isoform 1 of GSN |
| P17742 | PPIA |
| P20029 | HSPA5 |
| P27773 | PDIA3 |
| P62962 | PFN1 |
| Q05793 | HSPG2 |
| Q61001 | LAMA5 |
| Q61292 | LAMB2 |
| Q8VDD5 | MYH9 |
| Q9CYL5 | GLIPR2 |

**Table S5 -** Extracellular protein abundance in EPIC IM, EPIC SM, Matrigel and DMEH. The relative abundance of the top 5 extracellular proteins in the total pool of proteins identified per condition is shown. Relative abundances were calculated taking into acount each protein abundance from each sample. Abundance was averaged for each protein. Then, averaged abundances were summed. Finally, each averaged abundance was divided by the total sum to obtain in relative abundance. This was done for each sample type (EPIC IM, SM, DMEH, and Matrigel).

| **EPIC IM** | | **EPIC SM** | |
|---|---|---|---|
| **Protein** | **Relative abundance (%)** | **Protein** | **Relative abundance (%)** |
| HTRA1 | 15.21 | FN | 40.43 |
| SDF1 | 5.26 | MYH9 | 19.76 |
| FN | 3.36 | HSP90B1 | 2.92 |
| EMILIN-1 | 3.01 | ANXA2 | 2.55 |
| LAMA1 | 2.76 | HSPA5 | 1.94 |
| **E17.5 decellularized hearts** | | **Matrigel** | |
| **Protein** | **Relative abundance (%)** | **Protein** | **Relative abundance (%)** |
| FN | 0.82 | LAMA1 | 33.09 |
| HSPG2 | 0.61 | LAMB1 | 22.30 |
| HSPA5 | 0.28 | LAMC1 | 19.43 |
| COL1A2 | 0.27 | NID-1 | 14.55 |
| MYH9 | 0.23 | HSPG2 | 1.62 |
